# Soft, dynamic hydrogel confinement improves kidney organoid lumen morphology and reduces epithelial*–*mesenchymal transition in culture

**DOI:** 10.1101/2021.11.10.467741

**Authors:** Floor A.A. Ruiter, Francis L.C. Morgan, Nadia Roumans, Anika Schumacher, Gisela G. Slaats, Lorenzo Moroni, Vanessa L.S. LaPointe, Matthew B. Baker

**Affiliations:** MERLN Institute for Technology-Inspired Regenerative Medicine, Department of Complex Tissue Engineering, Maastricht University, Universiteitssingel 40, 6229 ER Maastricht, the Netherlands; MERLN Institute for Technology-Inspired Regenerative Medicine, Department of Cell Biology–Inspired Tissue Engineering, Maastricht University, Universiteitssingel 40, 6229 ER Maastricht, the Netherlands; Department II of Internal Medicine and Center for Molecular Medicine Cologne, University of Cologne, Faculty of Medicine and University Hospital Cologne, Cologne, Germany, Cologne Excellence Cluster on Cellular Stress Responses in Aging-Associated Diseases (CECAD), University of Cologne, Faculty of Medicine and University Hospital Cologne, Cologne, Germany

**Keywords:** Kidney organoids, Dynamic hydrogels, Viscoelastic, Epithelial-Mesenchymal Transition, Primary cilia

## Abstract

Pluripotent stem cell–derived kidney organoids offer a promising solution to renal failure, yet current organoid protocols often lead to off-target cells and phenotypic alterations, preventing maturity. Here, we created various dynamic hydrogel architectures, conferring a controlled and biomimetic environment for organoid encapsulation. We investigated how hydrogel stiffness and stress relaxation affect renal phenotype and undesired fibrotic markers. We observed stiff hydrogel encapsulation led to an absence of certain renal cell types and signs of an epithelial– mesenchymal transition (EMT), whereas encapsulation in soft-stress-relaxing hydrogels led to all major renal segments, fewer fibrosis/EMT associated proteins, apical proximal tubule enrichment, and primary cilia formation, representing a significant improvement over current approaches to culture kidney organoids. Our findings show that engineering hydrogel mechanics and dynamics has a decided benefit for organoid culture. These structure–property– function relationships can enable rational design of materials, bringing us closer to functional engraftments and disease-modelling applications.

## MAIN

Within the organoid field, various hydrogels have been investigated to influence cell behaviour^1, 2^, including biopolymer-based hydrogels (e.g. alginate)^3^, fully synthetic materials (polyethylene glycol (PEG) and polyacrylamide)^4, 5^, and bio hybrids (Matrigel combined with PEG, fibrin, or alginate)^6^. However, these hydrogels were mainly used in intestinal^7^, pancreatic^8^, neural^9^ and hepatic^10^ organoid cultures, while few have been applied in kidney organoid culture^5, 11^. Moreover, existing synthetic hydrogels for organoid culture largely rely on covalent or non-reversible cross-linking interactions, while the natural extracellular matrix (ECM) is dynamic^12^. Dynamic covalent cross-linked hydrogels have been of increased interest in the biomaterials field, as they allow recapitulation of both stiffness and dynamic stress-relaxing character of the native ECM^12,13,14,15,16^. To date, dynamic hydrogels have been observed to influence cell fate when single cells were encapsulated, as observed with extended motor neurons axon bodies^17^, 3D cell spreading and focal adhesion of hMSCs^18^, and increased cartilage matrix formation by chondrocytes^19^, but their application in aggregate or organoid culture is less studied.

Kidney organoids derived from induced pluripotent stem cells (iPSCs) mimic the organogenesis of the human kidney^20, 21^. In addition to their potential for studying development, disease modelling, and drug screening, kidney organoids can be transplanted as a functional graft^22, 23^ in patients with chronic kidney disease (CKD), which affects 11–13% of the population worldwide^24^. Nevertheless, there are still many challenges to overcome before organoids are suitable for widespread clinical application. For example, current kidney organoids resemble an immature developing kidney at both the transcriptional^25^ and morphological level,^26^ and prolonged culture does not improve their maturation. Moreover, morphological changes, an upregulation of off-target cell populations^25^ and aberrant ECM, containing increased types I and VI collagen and alpha smooth muscle actin (aSMA), are observed^11^. We have previously shown a reduction in the onset of the aberrant ECM expression and off-target cell populations by encapsulating the organoids in a soft thiol-ene cross-linked alginate hydrogel^11^. This initial study showed the importance of the surrounding environment on the organoid ECM, which led us to wonder how the mechanical stiffness and hydrogel dynamics of the encapsulating matrix can affect organoids.

Here, we tested the influence of dynamic hydrogels on kidney organoids by designing a small hydrogel library based on oxidised alginate: three hydrogels of tuneable stiffness (ranging from 0.1 to 20 kPa), and two soft hydrogels (0.1 kPa) with different stress relaxation (slow and fast). We used an imine-type dynamic covalent cross-linking possessing a range of equilibrium constants (*K*_eq_) that affect the hydrogel stiffness and tuneable hydrolysis rates (*k*_-1_) that change the rate of cross-link rearrangement and stress relaxation^27^. Kidney organoids cultured until day 7+14 (7 days of iPSC differentiation and 14 days of organoid culture on an air–liquid interface) were encapsulated in these hydrogels and cultured for 4 subsequent days (**Figure 1a**). We confirmed the formation of renal structures by immunohistochemistry. We investigated the expression of off-target ECM and EMT-related markers, as well as variations in the formation of lumen and cilia structures in renal organoids cultured in hydrogels of different stiffness and stress relaxation. Soft hydrogels with fast-relaxation properties resulted in more mature kidney organoids, determined by the above parameters, and compared to the stiffer hydrogels or slow-relaxing hydrogels. Our findings reinforce the concept of carefully selecting not only the stiffness, but also stress relaxation properties of encapsulating matrices and highlight the potential of tuning hydrogel properties to influence organoid cultures.

**Figure 1.**
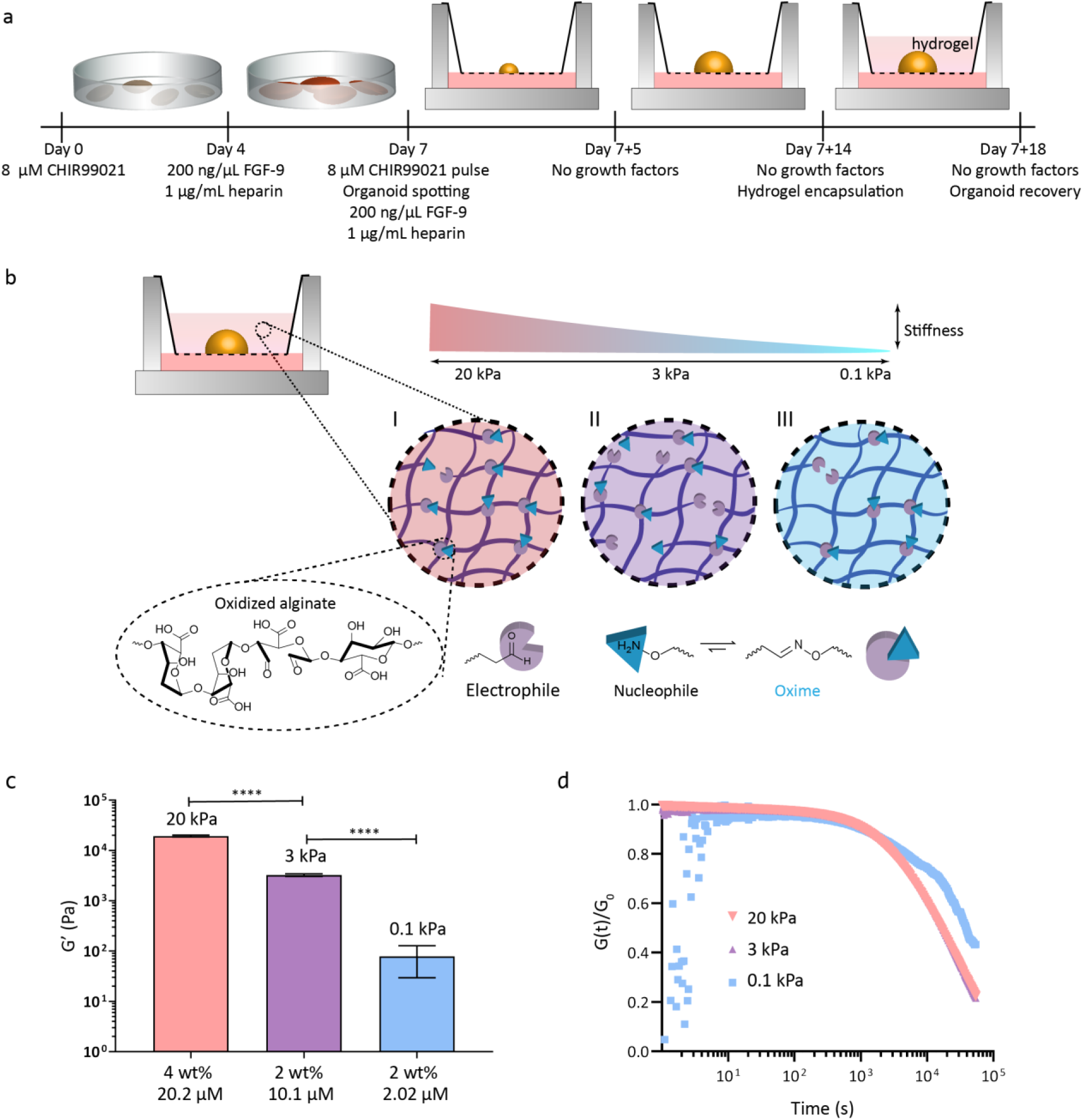
Hydrogels of varying stiffness were designed. (a) Overview of iPSC differentiation and organoid generation. iPSCs were differentiated for 7 d, after which they were grown and matured as organoids for 14 d on the air–liquid interface (day 7+14). Organoids were then encapsulated in the different hydrogels (or left on the air–liquid interface as a control) and cultured for 4 subsequent d until day 7+18. (b) Schematic of the different hydrogel systems. All hydrogels were based on oxidised alginate cross-linked with oxime. Increasing the percentage (by weight) of oxidised alginate resulted in increased entanglement and binding sites, and therefore increased stiffness. Similarly, an increase in oxime increased the cross-linking density, and therefore increased the stiffness of the hydrogel. (c) The shear moduli (G’) of the three different hydrogel compositions (N=3, error bars representing standard deviation), of 20 kPa, 3 kPa and 0.1 kPa for the 4% alginate–20.2 μM oxime; 2% alginate–10.1 μM oxime, and 2% alginate–2.02 μM oxime hydrogels, respectively. The hydrogels had significantly different stiffnesses (one-way ANOVA, *p* < 0.0001). (d) The stress relaxation of the different hydrogels compositions was similar (one-way ANOVA, *p*=0.32), with t_1/2_ values from 1.6–3.9·10^4^ s, despite their different stiffness.

### Hydrogels were designed with varying stiffness

To determine the role of hydrogel stiffness on renal organoid phenotype and ECM deposition, we designed three alginate hydrogels with varying stiffness (**Figure 1b**). We selected sodium alginate, a naturally-derived, biocompatible, and non-adhesive biomaterial, which is biodegradable when oxidised and has been used in FDA-approved applications^28^. Sodium alginate can form an ECM-like hydrogel and has previously been shown to support culture of kidney organoids^11^. We oxidised the alginate to obtain aldehyde groups for imine-type cross-linking, which allows for dynamic reshuffling of the cross-links in cell culture conditions^27^.

Sodium alginate was oxidised using sodium periodate (NaIO_4_) at a 10% theoretical degree of oxidation, then characterised by ^1^H-NMR (**Figure S1**) and gel permeation chromatography (GPC; **Figure S2** and **Table S1**). As a cross-linker, we used a small bifunctional oxime (O,O’-1,3-propanediylbishydroxylamine). Starting at 2 wt% oxidised alginate, increasing the concentration of bifunctional oxime increased the cross-linking density, which subsequently resulted in an increased stiffness, changing from 0.1–3.0 kPa for 2.02 to 10.1 µM of oxime cross-linker, respectively **(Figure 1b–c** and **S3a–c)**. To obtain a higher stiffness, we increased the weight percentage of the oxidised alginate (from 2 to 4 wt%) while keeping the 1:1 oxime to aldehyde ratio (20.2 µM oxime cross-linker), which resulted in a hydrogel with a stiffness of 20 kPa **(Figure 1c** and **S3a–c)**. While the stiffness of the hydrogels was significantly different in each composition (*p*<0.0001), the characteristic stress relaxation times (t_1/2_, defined as the time taken for the relaxation modulus to reach half of its initial value) were similarly long, with values from 1.6–3.9·10^4^ s (*p* = 0.32, **Figure 1d)**

### Kidney organoids formed in all hydrogels but selective renal cell types were absent in the 20 kPa hydrogel

Kidney organoids were cultured until day 7+14, at which point they were encapsulated in the different hydrogels for 4 additional days of culture **(Figure 1a)**. The starting point of day 7+14 was selected because our previous work demonstrated that an overexpression of type 1a1 collagen began at day 7+14 of organoid culture and could be significantly reduced by encapsulating the organoids in a hydrogels for 4 additional days^11^. The hydrogel solutions were added on top of the organoid on the air–liquid interface, and hydrogel cross-linking was observed after approximately1 h incubation at 37°C. After 4 d (day 7+18), the hydrogels were removed, and organoids were stained with calcein AM and EthD-1 to assess live/dead cells (**Figure S3e**). We detected no difference in the morphology of the organoids (bright-field microscopy, **Figure S4a)** or live/dead cells compared to organoids cultured on the air–liquid interface at day 7+18 **(Figure S4b)**.

To determine the effect of the hydrogel stiffness on the presence of different renal segments, the organoids were stained with relevant markers for distal tubules (E-cadherin; ECAD), glomeruli (nephrin; NPHS1), interstitial cells (homeobox protein Meis 1/2/3; MEIS1/2/3), loop of Henle (NKCC2; SLC12A1), and proximal tubules (LTL). The organoids recovered from the 0.1 and 3 kPa hydrogel possessed all expected segments (**Figure 2c-d** and **Figure S5c-d**) and had no distinguishable differences from organoids cultured on the air–liquid interface (**Figure 2a** and **Figure S5a**). However, organoids encapsulated in the stiffer 20 kPa hydrogel lacked interstitial and loop of Henle cells, and showed diminished lumen structures (LTL) and distal tubules (ECAD) (**Figure 2b and S5b**) compared to organoids cultured on both the air–liquid interface and other hydrogels (0.1 and 3 kPa).

**Figure 2.**
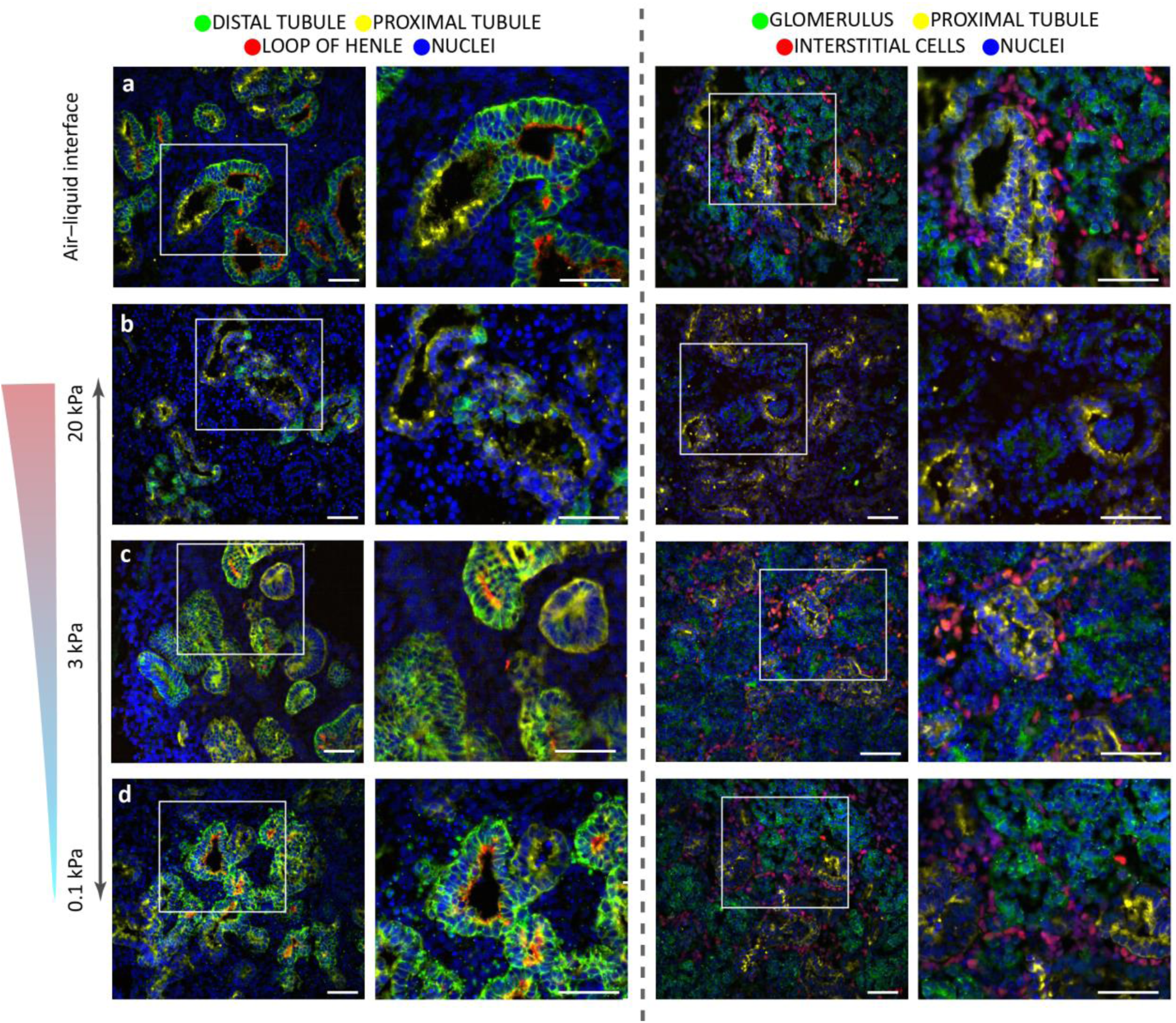
Kidney organoids formed in all dynamic hydrogels but renal cell types were absent in the 20 kPa hydrogel. Culture of the organoids for 4 days (from day 7+14 to 7+18) in hydrogels did not affect the presence of nephron segments in the 0.1 kPa (d) or 3 kPa (c) hydrogels when compared to organoids grown on an air–liquid interface until day 7+18 (a). Only the organoids encapsulated in the stiffest 20 kPa hydrogel (b) lacked loop of Henle segments (two left columns) and interstitial cells (two right columns). Staining for distal tubules (E-cadherin; ECAD); the loop of Henle (NKCC2; SLC12A1); proximal tubules (lotus tetragonolobus lectin; LTL); glomeruli (nephrin; NPHS1); and interstitial cells (homeobox protein Meis 1/2/3; MEIS1/2/3) as indicated. DAPI staining (blue) for nuclei. The white box denotes the area of interest enlarged in the respective right panel. Scale bars: 50 µm. Representative images of N=3 organoid batches with n=3 organoids per batch. See single channels in Figure S5.

### No epithelial–mesenchymal transition (EMT) observed in organoids encapsulated in the soft 0.1 kPa hydrogels

Because previous work demonstrated that prolonged (>7+14 d) organoid culture led to protein expression indicating fibrosis, namely collagens 1a1 and 6a1^11^, we wished to determine whether the kidney organoids encapsulated in the hydrogels showed fibrosic markers. Immunohistochemistry data showed a reduction of type 1a1 collagen (**Figure S6b–d**) in organoids encapsulated in all hydrogels compared to the air–liquid interface control (**Figure S6a**). In contrast, the expression of type 6a1 collagen was unchanged, indicating that the hydrogel stiffness had a selective modifying effect on the type 1a1 collagen. This selective effect has been observed previously when kidney organoids were encapsulated in a soft (0.2 kPa), thiol-ene cross-linked, alginate hydrogel^11^, and suggests a reduced fibrotic phenotype in organoids encapsulated in soft hydrogels.

Since EMT is an early marker of renal fibrosis^29, 30^, we also analysed the expression of EMT markers to determine the influence of hydrogel stiffness. We began by analysing single-cell RNA sequencing datasets from literature of kidney organoids culture up to day 7+27^25^ for EMT-related markers. We found the following indicators of an EMT: upregulation of twist family bHLH transcription factor 1 (*TWIST1*), snail zinc-finger transcriptional factor 1 (*SNAI1* encoding SNAIL), N-cadherin (*CDH2*), aSMA (*ACTA2*), and vimentin (*VIM*), and the downregulation of E-cadherin (*CDH1*) (**Figure 3a**). These EMT markers were deregulated from day 7+12 and found in all cells cultured at day 7+27 on the air–liquid interface, especially the mesenchymal marker vimentin, while the epithelial marker E-cadherin was downregulated (**Figure S7**).

**Figure 3.**
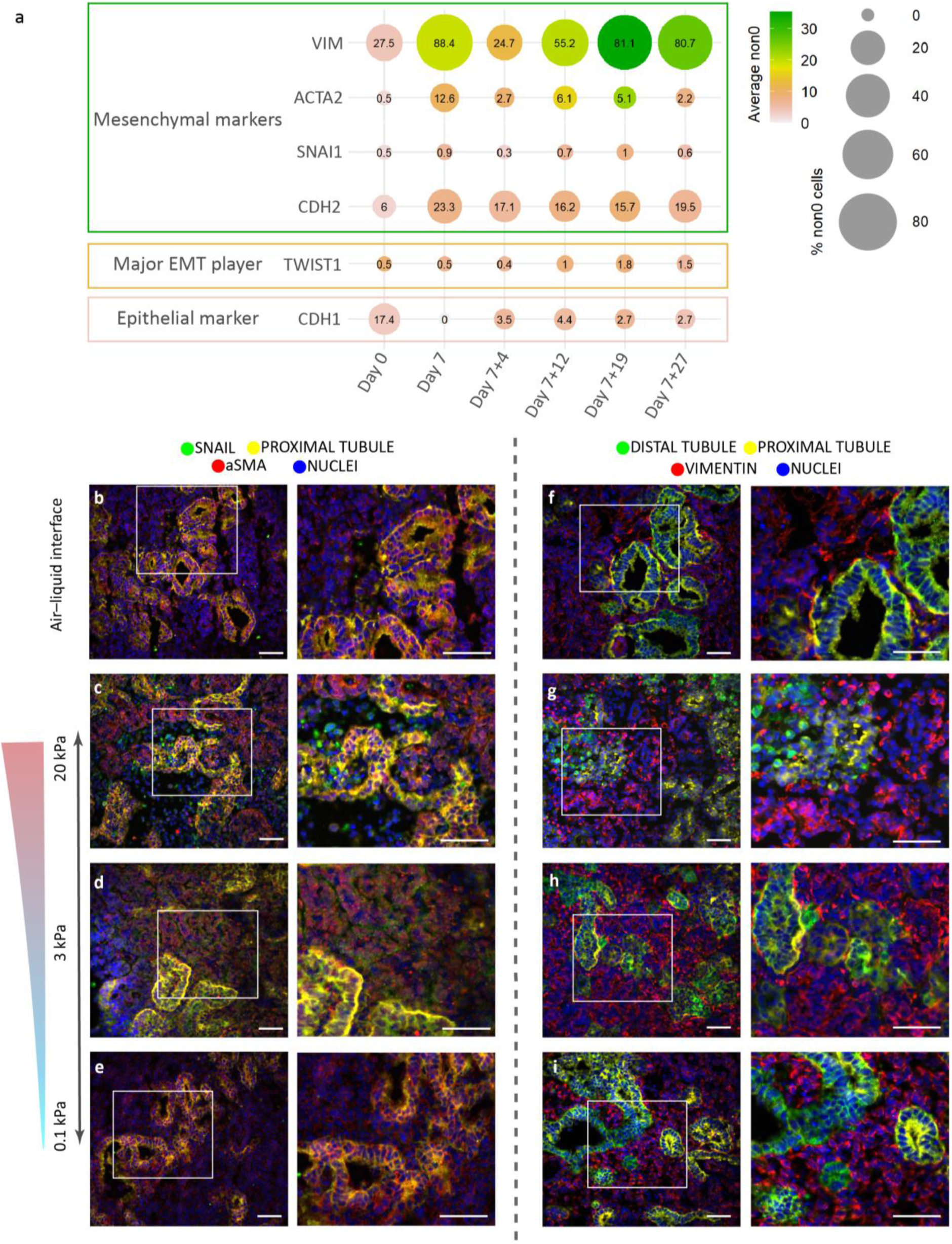
Evidence of epithelial–mesenchymal transition (EMT) observed in organoids encapsulated in the 3 and 20 kPa hydrogels. (a) Analysing single-cell RNA sequencing datasets from literature^25^ showed an increase of number of cells that expressed the EMT-related marker (*TWIST1*) and mesenchymal markers (*VIM, ACTA2, SNAI1,* and *CDH2*) and a reduced number of cells with the epithelial marker (*CDH1*). (b–i) Encapsulated organoids until day 7+18 showed an expression of the EMT transition marker SNAIL (green, two left columns) in the 20 kPa (c) and 3 kPa (d) hydrogels, whereas no expression was detected in the 0.1 kPa hydrogel (e) or the air–liquid interface control (b). Moreover, higher expression of the mesenchymal marker vimentin (red, two right columns) was observed in the 20 kPa (g) and 3 kPa (h) hydrogels compared to the air–liquid interface (f). Staining for distal tubules (E-cadherin: ECAD), proximal tubules (lotus tetragonolobus lectin: LTL), and fibroblasts (alpha smooth muscle actin: αSMA) as indicated. DAPI staining (blue) for nuclei. The white box denotes the area of interest enlarged in the respective right panel. Scale bars: 50 µm. Representative images of N=3 organoid batches with n=3 organoids per batch. See single channels in Figure S7.

For organoids cultured in the hydrogels up to 7+18 d, immunohistochemistry showed the classic EMT markers in the 3 and 20 kPa hydrogels. SNAIL-positive cells were found in organoids cultured in the 3 kPa (**Figure 3d**) and 20 kPa (**Figure 3c**) hydrogels, but not in the 0.1 kPa hydrogel (**Figure 3e**) or air–liquid interface (**Figure 3b**). Organoids cultured in the two stiffer hydrogels (3 and 20 kPa; **Figure 3d and c**) expressed more aSMA than organoids cultured in the 0.1 kPa hydrogel (**Figure 3e**) and the air–liquid interface (**Figure 3b**). Expression of the mesenchymal marker vimentin was observed in all organoids (**Figure 3f–i**), with higher expression in the organoids in the 3 and 20 kPa hydrogels (**Figure 3h and g**). Vimentin expression establishes the presence of mesenchymal cells and is required for EMT-related renal fibrosis^30,31,32^, where it is upregulated in proximal tubular cells when repair is initiated after epithelial injury^33^. In all organoids, vimentin did not co-stain with the proximal tubule and distal tubule cells, but was located between those segments **(Figure 3f–i and S8f-i)**. These findings, together with the absence of interstitial (MEIS1/2/3^+^) cells, (**Figure 2b**), increased aSMA-positive cells (**Figure 3c**), reduced lumen structures (**Figure S5b and S9c**), and diminished distal tubules (**Figure 2b**) in the stiffest 20 kPa hydrogel, suggest some epithelial cells have turned to mesenchymal cells and are contributing to the organoid stroma.

Our observations of fibrogenesis in organoids cultured on stiffer hydrogels is consistent with the literature reporting a link between fibrosis and tissue stiffness. For example, the kidney becomes stiffer (>15 kPa) with increasing fibrosis^34, 35^. The stiff hydrogel of >20 kPa in this study may activate the EMT pathways leading to fibrosis. Ondeck et al.^36^ saw a similar trend of increased EMT markers TWIST and SMAD2/3 in mammalian epithelial cells when dynamically increasing the stiffness of a methacrylate glycosaminoglycan hyaluronic acid hydrogel from 0.1 kPa to 3000 kPa. Others have reported that a softer environment can accelerate the differentiation of iPSC derived kidney organoids^5^ and can prime undifferentiated cells to lineage commitment^37^. Moreover, Chen et al.^38^ observed the prevention of TGF-β1– induced EMT when porcine kidney proximal tubule cells were cultured on collagen type 1– coated polyacrylamide gels of ∼0.2 kPa stiffness, while cells on a stiffer matrix (>0.7 kPa) highly expressed mesenchymal markers. Indeed, we observed the EMT marker SNAIL in the 3 kPa hydrogel but not in the 0.1 kPa hydrogel. Softer substrates are also more biologically relevant for a developing kidney: our organoids biologically resemble an embryonic kidney^11^, which has a stiffness of <1 kPa^39^, while our 3 kPa hydrogel is similar to the stiffness of an adult kidney of 5–10 kPa^35^. These finding combined show increased evidence that a softer hydrogel correlates with better performance of organoids and reduced expression of EMT markers.

### A fast-relaxing dynamic hydrogel further reduced the undesired EMT-related marker expression pattern

Beside stiffness (expressed as the shear moduli), stress relaxation plays a role in the cellular response to its surrounding material^18, 27, 40,41,42^. We had so far kept the stress relaxation of the hydrogel similar **(Figure 1d)** to compare only the effect of stiffness **(Figure 1c)**. To examine the effect of stress relaxation, we designed a fourth hydrogel with a similar stiffness as the soft oxime cross-linked hydrogel (∼0.1 kPa, **Figure 4a–b**) but with a faster stress relaxation time by using hydrazone as the cross-linker (**Figure 4a**). The imine group hydrolysis rate (*k*_-1_) can be tuned by changing the electronegativity of the alpha group to the primary amine^27^. In this case, we decreased the electronegativity of the alpha group by using hydrazone (adipic dihydrazide) instead of oxime to form a faster stress-relaxing hydrogel (**Figure 4a**). A higher concentration of the cross-linker hydrazone (10.1 µM) with 2% oxidised alginate was used to obtain a hydrogel of similar stiffness as the soft oxime hydrogel of 0.1 kPa (**Figure 4b and S10a-b**). The characteristic stress relaxation time of the resulting hydrazone hydrogel is an order of magnitude faster, with a t_1/2_ value of 4.1·10^3^ s, compared to the 3.9·10^4^ s for the soft 0.1 kPa oxime hydrogel (**Figure 4c)**.

**Figure 4.**
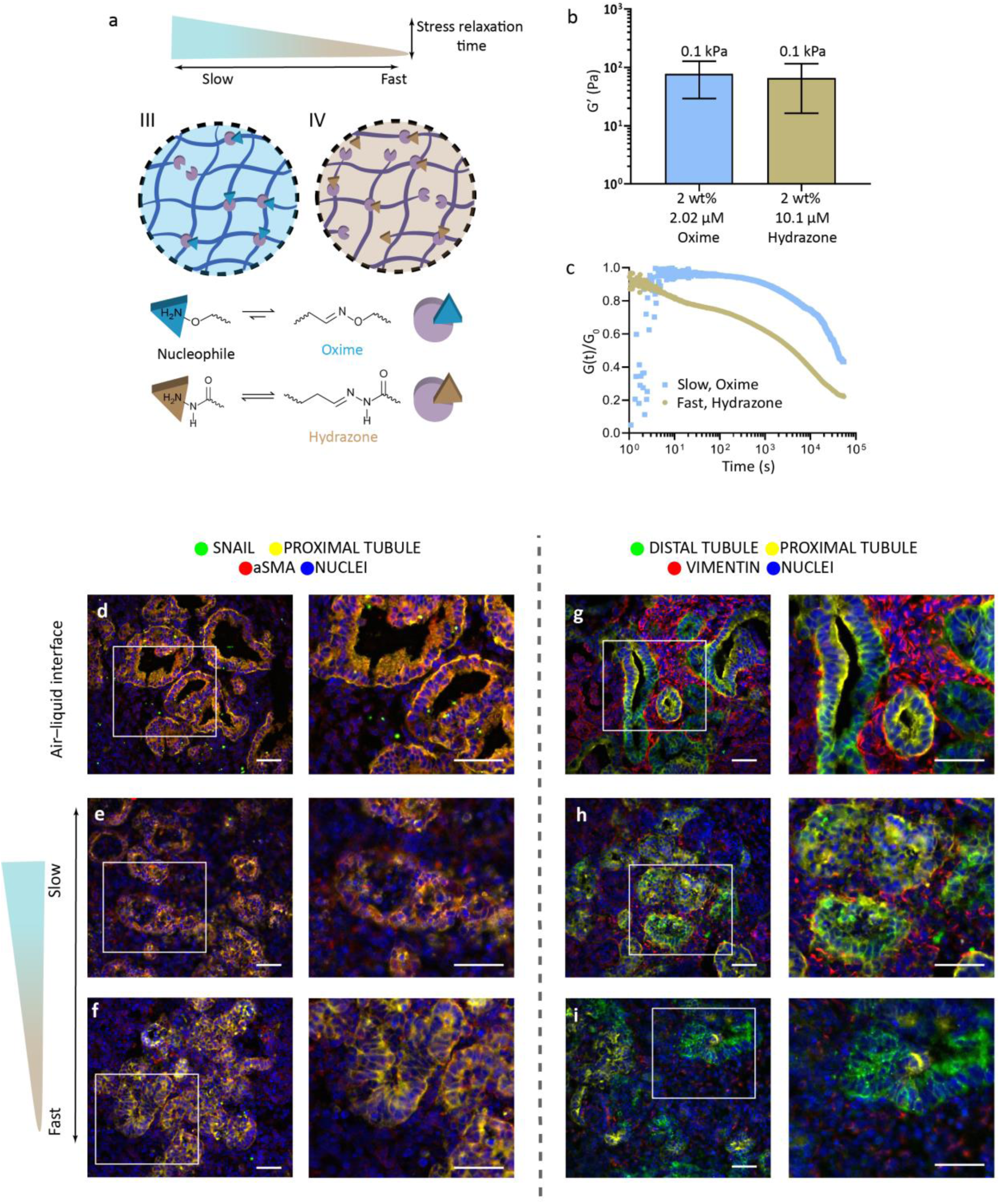
Fast-relaxing dynamic hydrogel further reduced the undesired EMT-related phenotype. (a) Schematic of two alginate hydrogels with different cross-linkers—hydrazone or oxime—to influence stress relaxation times. (b) The stiffness of both hydrogels is approximately 0.1 kPa (0.08 ± 0.05 and 0.07 ± 0.05 kPa for 2.02 μM oxime and 10.1 μM hydrazone, respectively; error bars represent the mean ± SD, t-test, *p*<0.0001). (c) The stress relaxation time of the fast-relaxing hydrazone hydrogel is an order of magnitude faster, with a value of 4.1·10^3^ s compared to the 3.9·10^4^ s for the soft 0.1 kPa oxime hydrogel (d–i). Encapsulated organoids until day 7+18 in the fast stress-relaxing hydrogel (f, i) showed less staining for aSMA (red, two left columns) and vimentin (red, two right columns) compared to those grown in the slow-relaxing hydrogel (e, h) and on the air–liquid interface (d, g). Staining for distal tubules (E-cadherin: ECAD), proximal tubules (lotus tetragonolobus lectin: LTL), and the mesenchymal markers (Zinc finger protein SNAI1: SNAIL and Vimentin) as indicated. DAPI staining (blue) for nuclei. The white box denotes the area of interest enlarged in the respective right panel. Scale bars: 50 µm. Representative images of N=3 organoid batches with n=3 organoids per batch. See single channels in Figure S12.

Organoids were encapsulated in the soft and faster stress-relaxing hydrogel at day 7+14 of culture, and were cultured to day 7+18. We detected no difference in the morphology of the organoids (bright-field imaging, **Figure S10d)** or ratio of live/dead cells compared to organoids cultured on the air–liquid interface **(Figure S10e)**. Recovered organoids showed all nephron segments similar to organoids cultured on the air–liquid interface (**Figure S11a-d).** A higher decrease of type 1a1 collagen expression was observed in organoids cultured in the soft, fast-relaxing hydrazone hydrogel (**Figure S11e**) compared to those cultured in the soft, slow-relaxing oxime hydrogel (**Figure S6a**), while no change in type 6a1 collagen staining was observed (**Figure S11f**) compared to culture on the air–liquid interface (**Figure S6a**).

Interestingly, there was less aSMA,–indicating the presence of myofibroblasts, and no EMT- related SNAIL marker – in the organoids encapsulated in the fast-relaxing hydrazone hydrogels (**Figure 4f**) compared to both those in the slow-relaxing, oxime cross-linked hydrogel (**Figure 4e**) and on the air–liquid interface (**Figure 4d**). Moreover, less expression of the mesenchymal marker vimentin was observed in the organoids encapsulated in the fast-relaxing hydrogel (**Figure 4i**) compared to the slow stress-relaxing (**Figure 4h**), and air–liquid interface (**Figure 4g**).

### Soft, fast-relaxing hydrogels result in apical LTL enrichment and increased ciliary length and frequency

Differences in lumen structure of the organoids cultured on the different hydrogels were observed, which we attribute to the stiffness and stress relaxation characteristics of the hydrogels. A significant increase of apical enrichment was observed in the LTL^+^ lumen of the organoids cultured in the two soft hydrogels (**Figure 5f-k-p-u-v and S9g-j**), while no clear lumina were observed in the stiffest hydrogels (20 kPa, **Figure 5b-h-m-r and S9c-d**). This indicated a link between stiffness and stress relaxation on the lumen maturation, in which the softer, fast-relaxing hydrogels increased the maturity.

**Figure 5.**
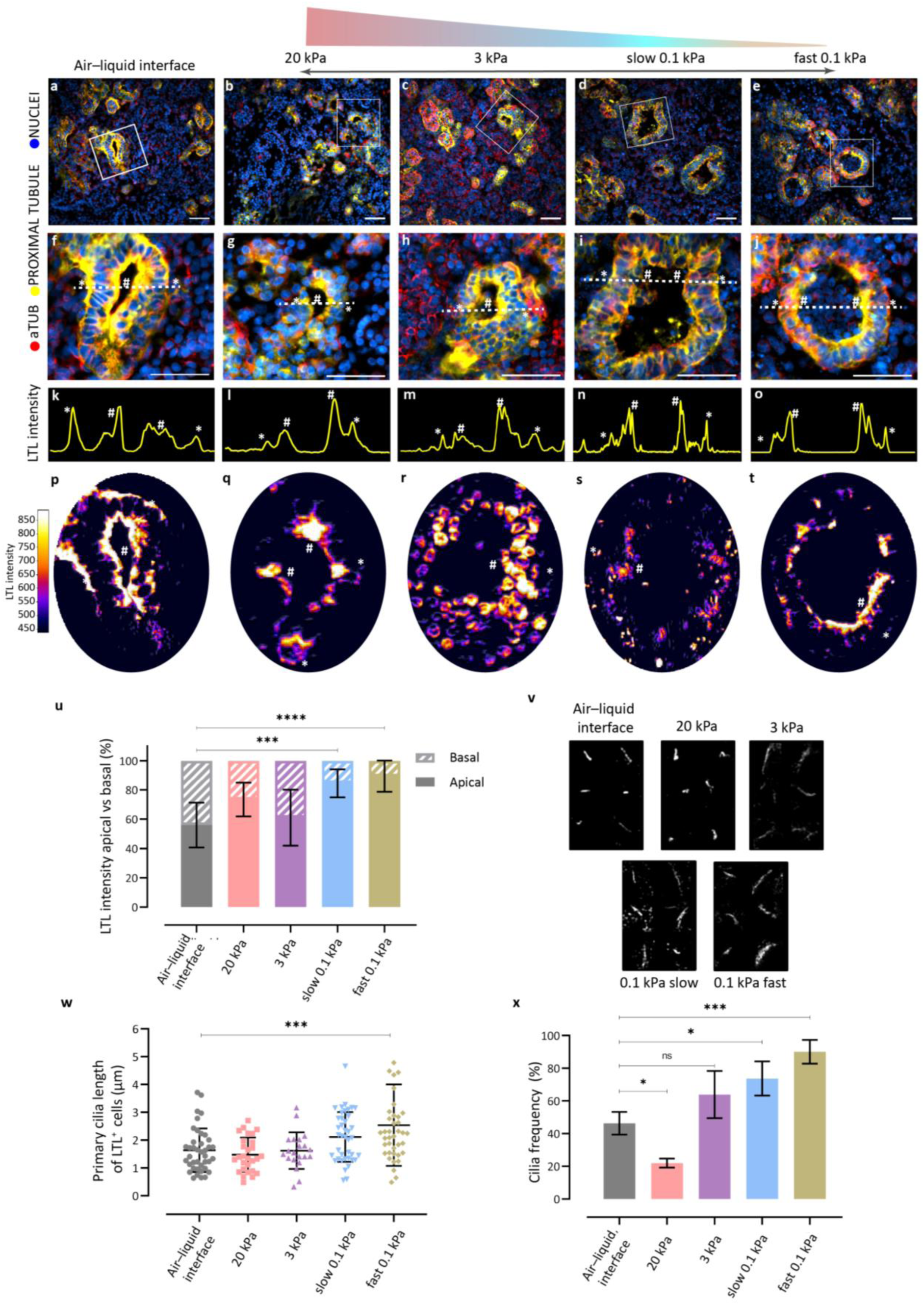
Stiffness and stress relaxation affect the polarity of the tubule lumen structures, ciliary length and frequency. Organoids encapsulated until 7+18 d in 0.1 kPa hydrogels (d,i,n,s and e,j,o,t) showed apical enrichment of LTL^+^ tubules compared to organoids grown in the other hydrogels. The air–liquid interface grown organoids show a clear apical and basal orientated LTL^+^ staining (a.f,k,p). When comparing LTL intensity apical vs basal a significant increase expression (***=0.0002 and ****<0.0001 for slow and fast-relaxing hydrogel respectively; one-way ANOVA) was observed in the soft hydrogels. Intensity plots (k-o) show the LTL intensity distribution over the dotted lines. The heat maps show overall LTL staining distribution over complete tubule structure. DAPI staining (blue) for nuclei. The white box denotes the area of interest enlarged in the respective right panel. Scale bars: 50 μm. Representative images of N=3 organoid batches with n=3 organoids per batch. See single channels in Figure S9 and S13. v) Primary cilia length was observed to vary in length in the different hydrogels compared to the air–liquid interface. Representative images of N=3 organoid batches (Figure S13) with n=3 organoids per batch. w) When measuring cilia length for cells with positive LTL staining, we observed significantly longer primary cilia for the 0.1 kPa fast hydrogel (*\*\*\*<*0.0005; one-way ANOVA), compared to the air–liquid interface. N=3 organoid batches per condition, 5 images per batch, 40 cilia of LTL^+^ cells were measured per condition. x) Cilia frequency was altered in the different hydrogels compared to the air–liquid interface. Cilia deficiency was observed in the 20 kPa hydrogel, while significant increase in cilia was observed in the soft, slow and fast-relaxing hydrogels. N=3 organoid batches per condition, 3 images per condition (ns= non-significant, *<0.05, ***<0.001, one-way ANOVA).

We hypothesised a link between the observed lumen structures and the accumulated stress, which the organoids experience through confinement in the stiffer hydrogels. Primary cilia play an essential role in sensing environmental cues (e.g. mechanotransduction), planar cell polarity of epithelial cells^43^, lumen formation, and EMT/fibrosis after acute kidney injury^44,45,46^, in which many factors (such as, chemical or physical cues) can modulate the ciliary length and frequency. Therefore, we stained for primary cilia (acetylated α-tubulin (aTUB) and ADP Ribosylation Factor Like GTPase 13B (ARL13B)) to investigate whether ciliary frequency and length vary in the different hydrogels. High-resolution z-stack confocal images showed differences in ciliary length (**Figure 5v and S13)**. Significantly increased average ciliary length was observed in the softer hydrogels, 2.3 (*p*=0.0002) and 2.7 µm (*p*<0.0001) for slow and fast-relaxing 0.1 kPa, respectively, compared to 1.6 µm for the air–liquid interface (**Figure S13p**, all cell types). During kidney development, ciliary length in the nephron lumen significantly increases from 0.59 to 3.04 µm from renal vesicles to fetal nephron, respectively^45, 47^. When only comparing proximal tubular ciliary length, we observed a significant increase in length of 2.5 µm (*p*=0.0002) in the soft fast-relaxing hydrogel compared to 1.6 µm for the air–liquid interface (**Figure 5w**).

EMT can also trigger reduced ciliary frequency (deficiency) and length^48^. The stiffest hydrogel (20 kPa) resulted in a significant reduction of ciliary frequency (% of ciliated cells, *p*=0.029, **Figure 5x**), while 3 kPa does not show a significant difference. Both these hydrogels show EMT-related markers. In contrast, a significant increase in ciliary frequency was observed in the soft slow and fast relaxing hydrogels (*p*=0.015, *p*=0.0006, respectively, **Figure 5x**). This clearly shows that the organoids feel the mechanics of the confining hydrogel environment, in which a higher stiffness (20 kPa) results in reduction of cilia frequency and length, while softer hydrogels (0.1 kPa) significantly increase the frequency. However, only the fast-relaxing hydrogel shows a significant increase in cilia length, as well as frequency. This may be due to the fast stress-relaxation of the hydrogel network, which dissipates the tension build up, due to confinement as the organoid expand, throughout the hydrogel, reducing tension in the overall organoids. If we combine the significantly increased ciliary length and frequency with the clear reduction of mesenchymal cells (VIM), aSMA expression, and LTL apical enrichment in the organoids encapsulated in the soft fast-relaxing hydrogel, we can postulate a link between hydrogel stiffness and stress relaxation on lumen maturation. Confined in a hydrogel, the ability to disperse tension through rapid stress-relaxing results in a reduction of off-target cell types and increased lumen maturation.

### Outlook

The effect of hydrogel stiffness and stress relaxation has been well studied on individual cells, but fewer studies have reported their combined effects on aggregates and organoids. The potential to engineer a matrix that influences these large multicellular aggregates may have been overlooked due to the perception that relatively few cells interact with the matrix. As iterated in our findings, mimicking the dynamics of the ECM has repeatedly been shown to be important. Our studies indicate that the stiffness and stress relaxation of the surrounding environment had a direct impact on kidney organoids. Stiffer hydrogels caused an undesirable EMT and loss of lumen structures, while softer hydrogels positively reduced undesired ECM deposition, increased the maturity of lumen structures, and reduced mesenchymal cells. Future work will focus on specific tuning of the stress relaxation to mimic the developmental embryonic environment. Overall, our results show the importance and potential of tuning the hydrogel properties to influence kidney organoids. Properly engineered matrices for organoids could lead to a platform for culturing functional engraftments and to disease modelling applications by tuning the hydrogel to mimic pathological scenarios such as fibrogenesis.

## Acknowledgements

This work is supported by the partners of Regenerative Medicine Crossing Borders (RegMed XB), a public–private partnership that uses regenerative medicine strategies to cure common chronic diseases. It is financed by the Dutch Ministry of Economic Affairs by means of the PPP Allowance made available by the Top Sector Life Sciences & Health to stimulate public–private partnerships. MBB and VLSL would like to acknowledge funding from the Dutch Province of Limburg; FLCM, LM, and MBB would like to thank funding from NWO under project agreement 731.016.202 “DynAM”. The authors also would like to thank Christian Freund (hiPSC core facility, LUMC, Leiden, the Netherlands) for supplying the hiPSC line (LUMC0072iCTRL01) used in this study, and Nathalie Groen for conducting the scRNA sequencing analyses.

## Methods

### Oxidised alginate (Oxi-Alg) synthesis

Sodium alginate was oxidised (10%) as previously described.^27^ Briefly, purified sodium alginate (1.0 g, 1 equiv, 5.68 × 10^−3^ mol monomer, Manugel GMB, FMC, Lot No. G940200) was dissolved in 100 mL deionised H_2_O overnight. Sodium (meta)periodate (0.121 g, 0.1 equiv, 5.68 × 10^−4^ mol, Sigma-Aldrich) was added. The mixture was covered with aluminium foil and stirred in the dark for 17 h at room temperature (RT). The reaction was quenched by the addition of ethylene glycol (0.035 g, 0.1 equiv 5.68 × 10^−4^ mol, Sigma-Aldrich) and stirred for 1 h in the dark at RT. The resultant product was dialysed in a 10 kDa MWCO dialysis tube (Spectra/Por, regenerated cellulose, VWR) for 3 d in 100 mM, 50 mM, 25 mM, 12.5 mM and 0 mM NaCl in deionised H_2_O (changed twice daily), and was flash-frozen in liquid N_2_ and lyophilised. Oxidation was confirmed by ^1^H-NMR in deuterated water (D_2_O, **Figure S1**) by the appearance of the protons between 5.15 and 5.75 ppm, attributed to the formation of hemiacetal groups upon reaction of the aldehydes to neighbouring hydroxyl groups. Moreover, molecular weights of the product were determined via GPC (**Figure S2**).

### 1H-NMR analysis

^1^H-NMR spectra were recorded on a Bruker Avance III HD 700-MHz spectrometer equipped with a cryogenically cooled three-channel TCI probe in D_2_O with sodium trimethylsilylpropanesulfonate-d6 as an internal standard (DSS-d6, 2 × 10^-3^ M). Water suppression pulse sequence was applied to spectra. Spectra analyses were performed with MestReNova 11.0 software. Chemical shifts are reported in parts per million (ppm) relative to DSS-d6 (CH_2_, 0 ppm).

### GPC analysis

Agilent PEG calibration kit (PEG molecular weights up to 300,000 MW, Agilent Technologies) were used for calibration. The samples were dissolved at a concentration of 2 mg·mL^-1^ in 0.1 M NaNO_3_ H_2_O and filtered through 0.45 μm filters to remove any unwanted impurities. Samples MW were measured in aqueous 0.1 M NaNO_3_ eluent with a flow rate of 0.5 mL·min^-1^ at RT on a Prominence-I LC-2030C3D LC (Shimadzu Europa GmbH) and Shodex SB-803/SB-804 HQ columns (Showax Denko America, Inc). LabSolutions GPC software (Shimadzu Europa GmbH) was used to calculate the molecular weight and dispersity values (Table S1).

### Rheometry

Rheological characterisation on the hydrogels was performed on a DHR2 rheometer from TA Instruments. Time sweeps of preformed hydrogels were taken over 360 s with an 8 mm parallel plate geometry at 20 °C with an applied strain of 1% at 3.14 rad/s. During loading, the gap size was adjusted to achieve 0.1 N of normal force and varied between samples from 800 µm to 1150 µm with a mean of 1004 ± 76 µm. The shear storage modulus (G’) for a given sample was taken to be the mean recorded value with a minimum of three sample replicates per formulation (**Figure 1c and 5b**). Stress relaxation measurements were performed using a 20 mm cone-plate geometry equipped with solvent trap. Precursor solutions were mixed as described in **Table S2** to obtain the desired final concentrations of oxidised alginate and cross-linker. Following the final addition of the cross-linker, samples were vortexed for 10 s before immediately loading 80 µL into the rheometer. Time sweeps were measured over 3.5–9 h (maintained at 20 °C) to monitor crosslinking progress with an applied strain of 1% at 10 rad/s. Once a plateau was reached, a frequency sweep was performed from 100–0.1 rad/s with an applied strain of 1% and 10 pts/dec (**Figure S3c**). Finally, to measure the stress relaxation behaviour, the relaxation modulus was monitored over 15.5 h with an initial applied strain of 20% maintained over the course of the measurement (**Figure 1d and 5c**). Statistical analysis were performed in GraphPad 8.2.0 using one-way ANOVA.

### Swelling test of hydrogels under culture conditions

A swelling test to investigate real-world swelling under culture conditions was performed on transwell filters, as previously described^11^. Briefly, hydrogel solutions were prepared (**Table S1**). 500 µL of this solution was added onto the transwells with a 0.4 µm pore size (Corning, 12-well culture plate) without organoids and left to cross-link for 1 h. After which, STEMdiff APEL2 medium (1% (v/v) PFHM-II protein-free hybridoma medium (Thermo Fisher Scientific) and 1% (v/v) antibiotic/antimycotic (Thermo Fisher Scientific) was added below the transwells with hydrogels and incubated up to 98 h (37 °C). Transwells with hydrogels were weighted at 1, 2, 4, 6, 24, 48, 72, and 96 h. Hydrogel swelling ratio in % (*S_r_*) over time was calculated by: 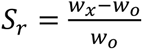, in which *w_0_* = initial hydrogel weight and *w_x_* = hydrogel weight at the *x* time point (**Figure S3d and S8d**). Graphad 8.2.0 software was used for statistical analysis using two-way ANOVA or unpaired t-test.

### Cell culture

As described previously^11^, hiPSC line LUMC0072iCTRL01 was generated from fibroblasts using the Simplicon RNA reprogramming kit (Millipore) by the hiPSC core facility at the Leiden University Medical Center. The cells were expanded in E8 medium (Thermo Fisher Scientific) on vitronectin-coated (0.5 μg·cm^-2^) plates and passaged with TrypLE Express (Thermo Fisher Scientific) twice weekly. For 24 h after each passage, cells were cultured in E8 medium supplemented with RevitaCell Supplement (Thermo Fisher Scientific). Subsequently, cells were cultured in E8 medium refreshed daily.

### Differentiation and organoid formation

Kidney organoids were produced from hiPSCs according to an established protocol (**Figure 1a**).^21^ Briefly, hiPSCs were seeded on 6-well plates with vitronectin-coating (0.5 μg·cm^-2^), at 7,300 cells per cm^2^ in E8 medium supplemented with RevitaCell Supplement. Differentiation was started after 24 h (day 0) by changing to STEMdiff APEL2 medium (STEMCELL Technologies) supplemented with CHIR99021 (8 µM, R&D Systems), 1% (*v/v*) antibiotic/antimycotic (Thermo Fisher Scientific), and 1% (*v/v*) PFHM-II protein-free hybridoma medium (Thermo Fisher Scientific). On day 4 of differentiation, the medium was changed to STEMdiff APEL medium supplemented with heparin (1 µg ml^-1^, Sigma-Aldrich) and FGF-9 (200 ng·ml^-1^, R&D Systems). By day 7, the cells formed a confluent monolayer and were subjected to a 1 h pulse of CHIR99021 (5 µM) in STEMdiff APEL medium before being harvested by trypsinisation. An aggregate of 500,000 cells was transferred to a Transwell filter with 0.4 µm pore size (Corning) in a 12-well culture plate to form the kidney organoids. Kidney organoids were cultured for 4 d (termed day 7+4) on an air–liquid interface with STEMdiff APEL2 medium supplemented with FGF-9 and heparin in the bottom compartment (450 µL) with media changes every 2 d. At day 7+5, STEMdiff APEL2 medium with no supplemented growth factors was used when media changes where performed every 2 d. The organoids were maintained until day 7+18, with or without hydrogel encapsulation from day 7+14 (**Figure 1a– b**).

### Hydrogel encapsulation

Oxidised alginate (120 mg, 10% oxidation, 6 wt% stock solution, UV sterilised, Oxi-alg) was dissolved in 2 mL STEMdiff APEL2 medium and left to dissolve overnight. *O*,*O*’-1,3- propanediylbishydroxylamine dihydrochloride (oxime, Sigma-Aldrich) or adipic dihydrazide (hydrazone, Sigma-Aldrich) stock solutions of 8 × 10^-2^ mol·L^-1^ were prepared in STEMdiff APEL2 medium (**Table S2**) and passed through a 0.2 µm sterilisation filter. The oxi-alg, crosslinker stock solutions, and STEMdiff APEL2 medium were added to obtain the desired hydrogel systems (**Table S2**) and vortexed before organoid encapsulation to form a 2% or 4% (w/v) sodium alginate solution. The hydrogel solutions were pipetted over the organoids on the top of the Transwell membrane at day 7+14 of culture (500 µL each). STEMdiff APEL2 medium (450 µL) was added to below the Transwell filters, and organoids encapsulated in the hydrogels were cultured for 4 additional days (until day 7+18, **Figure 1A-B**).

### Cryo-sectioning recovered organoids

At day 7+18, the hydrogels were removed and 4% paraformaldehyde (PFA) was added above and below the Transwell with the recovered organoid for 20 min at 4 °C. Organoids were cryo-sectioned as described previously^11^. Briefly, organoids were dehydrated overnight (PBS containing 15% (*w/v*) sucrose) at 4 °C followed by a second 2 d dehydration incubation (30% (*w/v*) sucrose). The dehydrated organoids were embedded in freezing solution (15% (w/v) sucrose and 7.5% (*w/v*) gelatin in PBS). The embedded organoids were placed in a beaker with isopentane and left to freeze in liquid N_2_ for several minutes. The frozen organoids were horizontally sectioned to 20 µm thickness at -18 °C.

### Immunohistochemistry

The embedding solution of the frozen organoid sections was removed by incubating for 15–20 min in PBS at 37 °C. The sections were washed (PBS), permeabilised (PBS with 0.5% (*v/v*) IGEPAL) for 15 min (RT), blocked (PBS with 5% (*w/v*) donkey serum, 1% BSA and 0.3 M glycine) for 20 min (RT), and incubated overnight at 4 °C (in the dark) with primary antibodies (PBS with 1% BSA and 0.3 M glycine) against: nephrin (NPHS1), lotus tetragonolobus lectin (LTL), Meis Homeobox 1/2/3 (MEIS1/2/3), E-cadherin (ECAD), solution carrier family 12 Member 1 (SLC12A1), type 1a1 and type 6a1 collagen, aSMA, vimentin, acetylated tubulin (a-tubulin), and Zinc finger protein SNAI1 (**Table. 1**). Subsequently, the slides were washed three times with PBS (1% BSA and 0.3 M glycine) and incubated with secondary antibodies including DAPI (0.1 µg/ml) for 1 h at RT in the dark: Alexa Fluor 488 (Thermo Fisher Scientific, 1:300, sheep/mouse/rabbit), Alexa Fluor 568 (Thermo Fisher Scientific, 1:300, mouse/rabbit), and Streptavidin Alexa Fluor 647 (Thermo Fisher Scientific, 1:100). Slides were washes three time in PBS and mounted with Mowiol mounting medium. Images were taken with an automated Nikon Eclipse Ti2-E microscope (20× or 40× air objective) or light microscope Leica TCS SP8 STED (100× objective).

**Table 1.** Primary antibodies used in this study.

| Primary Antigen | Mono- or Polyclonal | Source (catalog no.) | Species | Membrane or nucleus | Identifies | Dilution |
| --- | --- | --- | --- | --- | --- | --- |
| Acetylated α-Tubulin (aTUB) | Mono | MERCK (T7451) | Mouse | Membrane | Primary Cilia | (1:750) |
| aSMA | Mono | Sigma-Aldrich (A2547) | Mouse | Cytoplasm/ECM | Fibroblasts | (1:300) |
| ECAD | Mono | BD Biosciences (610181) | Mouse | Membrane | Distal tubule | (1:300) |
| LTL | - | Vectorlabs (B-1325) | Biotin conjugate | Membrane | Proximal tubule | (1:300) |
| MEIS1/2/3 | Mono | Santa-Cruz Biotechnology (SC-101850) | Mouse | Nucleus | Interstitial cells | (1:300) |
| NPHS1 | Poly | R&D Systems (AF426)9 | Sheep | Membrane | Podocytes | (1:300) |
| SLC12A1 | Poly | Thermo Fisher Scientific (HPA014967) | Rabbit | Membrane | Loop of Henle | (1:200) |
| SNAIL | Mono | Thermo Fisher Scientific (PA5-85493) | Rabbit | Nucleus | EMT transcription factor | (1:300) |
| Type I Collagen (COL1A1) | Mono | Abcam (ab6308) | Mouse | ECM | Type 1 Collagen | (1:300) |
| Type VI Collagen (COL6A1) | Mono | Genetex (GTX109963) | Rabbit | ECM | Type 6 Collagen | (1:300) |
| **Vimentin**<br>**(VIM)** | Mono | Thermo Fisher<br>Scientific<br>(MA3745) | Mouse | ECM | EMT ECM<br>marker | (1:300) |

### Cell viability assay

EthD1/calcein AM staining were used to determine cell viability. The encapsulating hydrogel and medium were removed. Recovered organoids on Transwells were incubated in EthD1 (4 µM) and calcein AM (2 µM) in PBS solution (top and bottom of the Transwell) for 30 min at RT. Organoids were imaged in PBS with an automated Nikon Eclipse Ti2-E microscope at 4×, 20×, or 40× air objective.

### Analysis of EMT-related gene expression in kidney organoids

Single-cell RNAseq data of iPSC-derived kidney organoids generated using the Takasato protocol^25^ were downloaded from the Gene Expression Omnibus (GEO: GSE118184) and analysed as previously described.^11^ Briefly, transcript count tables were analysed for each time point (day 0, 7, 7+5, 7+12, 7+19 and 7+27) by R software (3.6.2) and the Seurat package (version 3.2.0);with the exclusion of low-quality cells. The gene expression matrices were log-transformed using a scaling factor of 10,000 and normalised for sequencing depth per cell. Subsequently, the highest cell-to-cell variations were identified, scaled, and centred. These data were used for principal component analysis. Non-linear dimensional reduction was performed on selected principal components representing the true dimensionality of each dataset. Normalised markers gene expression of interest are presented in tSNE space.

### LTL polarity of Lumen structures

Immunohistochemistry images were processed in ImageJ. Plot profiles were analysed in ImageJ and heatmaps were generated by the interactive 3D surface plot plugin. Percentage of basal vs apical intensity were calculated by deducting plot profile from apical side from the full plot profile per lumen structure. (N=3 organoid batches, n=9 images per condition were processed). Statistical analysis were performed in GraphPad 8.2.0 using two-way ANOVA.

### Primary cilia length and frequency measurements

Z-stack confocal images measured (100× objective) were processed in ImageJ to Z projection. Length per cilia were measured using the straight-line function (N=3 organoid batches, n=5 images per condition were processed). Cilia frequency were measured using the cell counter plugin, cilia frequency were calculated by measuring the percentage of nuclei with primary cilia compared to the overall nuclei (N=3 organoid batches, n=3 images per condition were processed). Statistical analysis were performed in GraphPad 8.2.0 using two-way ANOVA.

## Additional information

The sugar monomers’ hydroxyl groups at C-2 and C-3 in the alginate chain were oxidised resulting in the breakage of the C–C bonds to form two aldehyde groups, which rapidly formed hemiacetals with neighbouring alcohol groups. These hemiacetal groups were confirmed via the appearance of the proton peaks at 5.2–5.8 ppm in the ^1^H-NMR spectra (**Figure S1**), confirming the oxidation. The molecular weight (MW) decreased from approximately 320 kDa in the starting material to 70 kDa due to the oxidisation breaking part of the backbone of the alginate structure (**Figure S2 and Table S1**).

## Supplementary figures and tables

**Figure S1.**
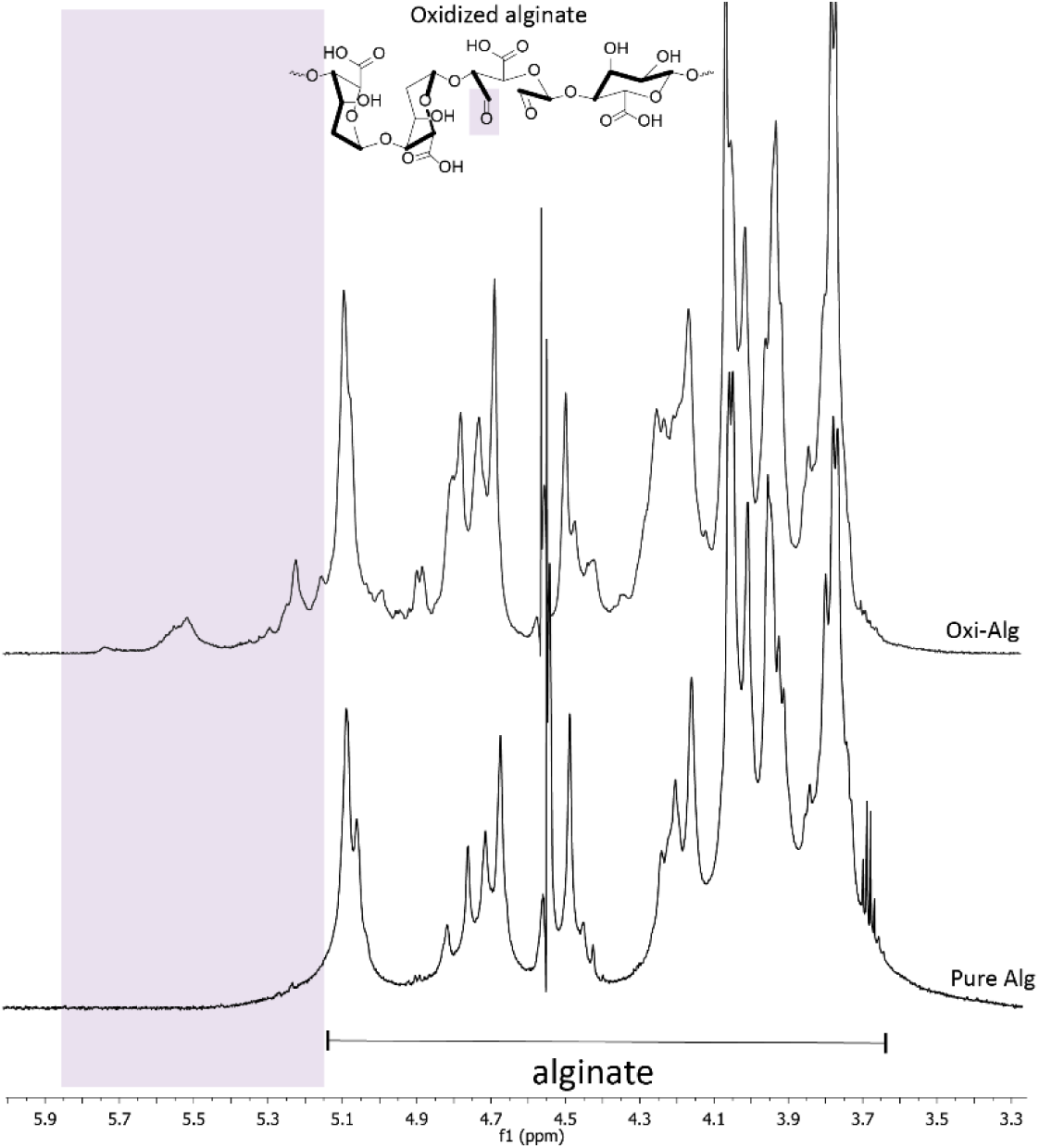
^1^H-NMR spectra of pure alginate (pure Alg; bottom) and oxidised alginate (oxi-alg; top). Oxidation of the alginate was confirmed by the appearance of the protons between 5.15–5.75 ppm, attributed to the formation of hemiacetal groups upon reaction of the aldehydes to neighboring hydroxyl groups (top spectra, in purple area), compared to the pure alginate NMR spectra, in D_2_O. DSS-d6 was used as internal standard and samples were measured at 325 K.

**Figure S2.**
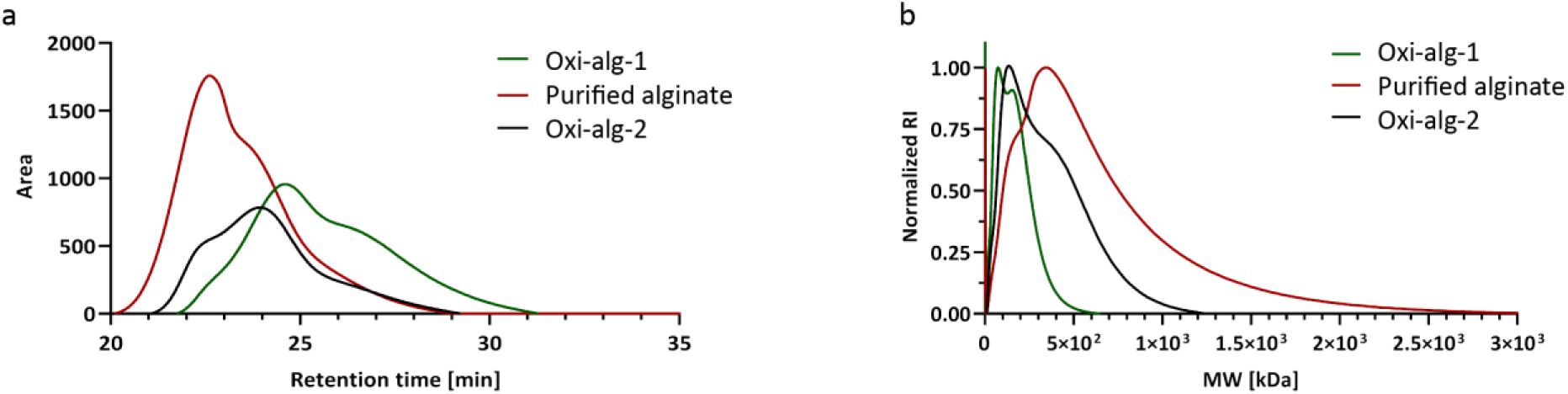
Reduced molecular weight confirms oxidation of the alginate. a) GPC retention time of the two oxidised alginate batches (oxi-alg-1 for the stiffness series and oxi-alg-2 for the fast-relaxing, 0.1 kPa hydrogel) compared to the pure alginate. b) Molecular weight of the oxidised alginates batches (oxi-alg-1 for the stiffness series and oxi-alg-2 for the fast-relaxing, 0.1 kPa hydrogel) compared to the pure alginate. As expected, the oxidation of the alginate backbone resulted in a reduction of the molecular weight of the resultant product; see values in Table S1.

**Figure S3.**
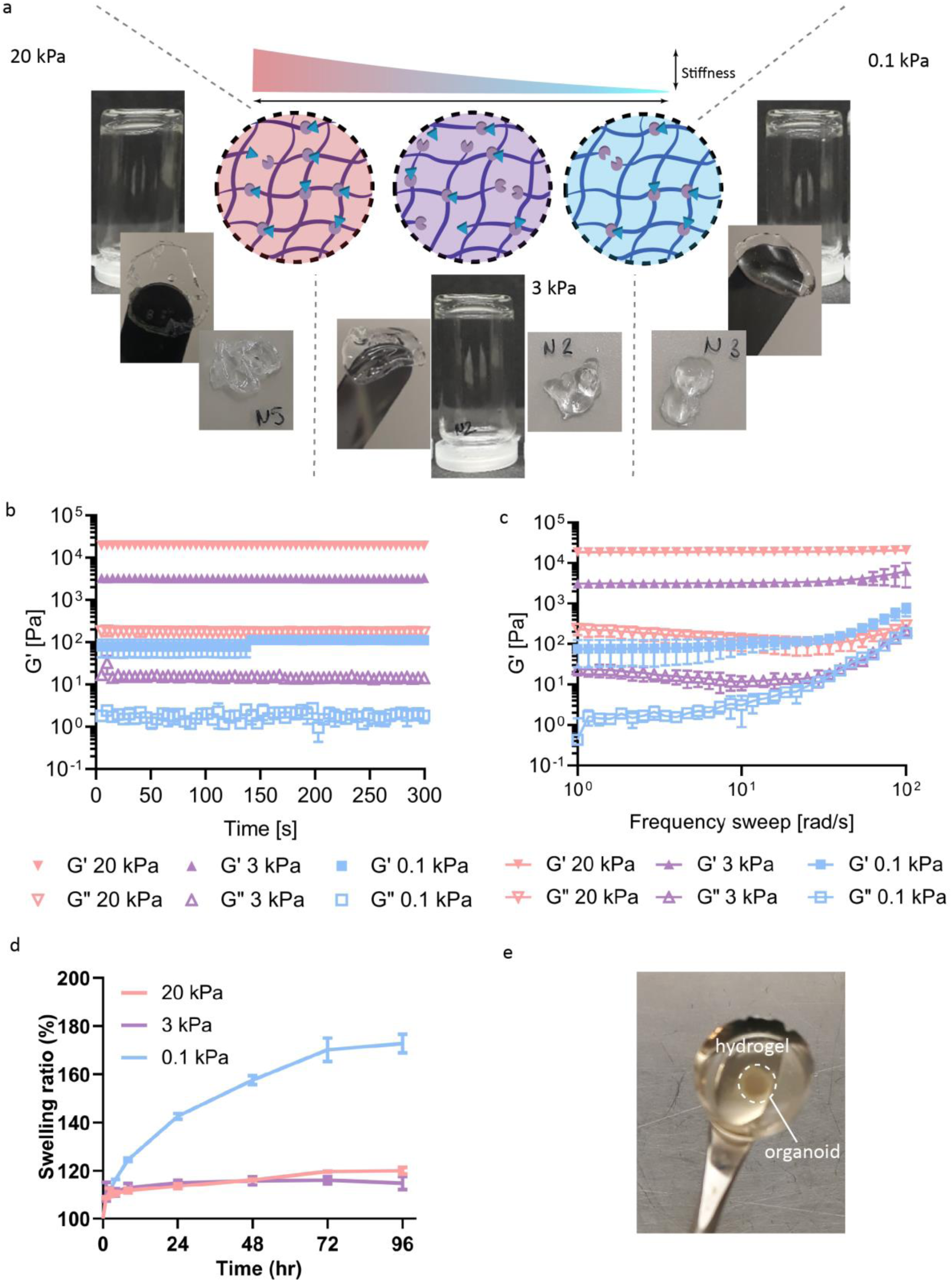
Hydrogel formation of different stiffnesses were confirmed and properties characterised. a) Hydrogel formation confirmed by inverted vial test (left image of each set); formed hydrogel on spatula (centre image of each set); and removed hydrogel from vial (right image of each set). The different systems were (left to right): 4% alginate–20.2 μM oxime (pink); 2% alginate–10.1 μM oxime (purple); 2% alginate–2.02 μM oxime (blue). b) Time sweep data of the three hydrogels was used to determine the stiffness values of the hydrogels (N=2, Figure 1C). Colour coding of hydrogels as indicated in panel a. c) Frequency sweep data of the hydrogels (N=2) showed they were frequency-independent. Colour coding of hydrogels as indicated in panel a. d) Swelling test showed a significant swelling of 172% for the 0.1 kPa hydrogel (two-way ANOVA, *p* < 0.0001, N=3) after 96 h incubation, compared to 120% swelling for the 3 kPa and 20 kPa hydrogels. Points and bars indicate mean ± standard error from three individual samples. e) Representative image of recovered organoid in the 20 kPa hydrogel.

**Figure S4.**
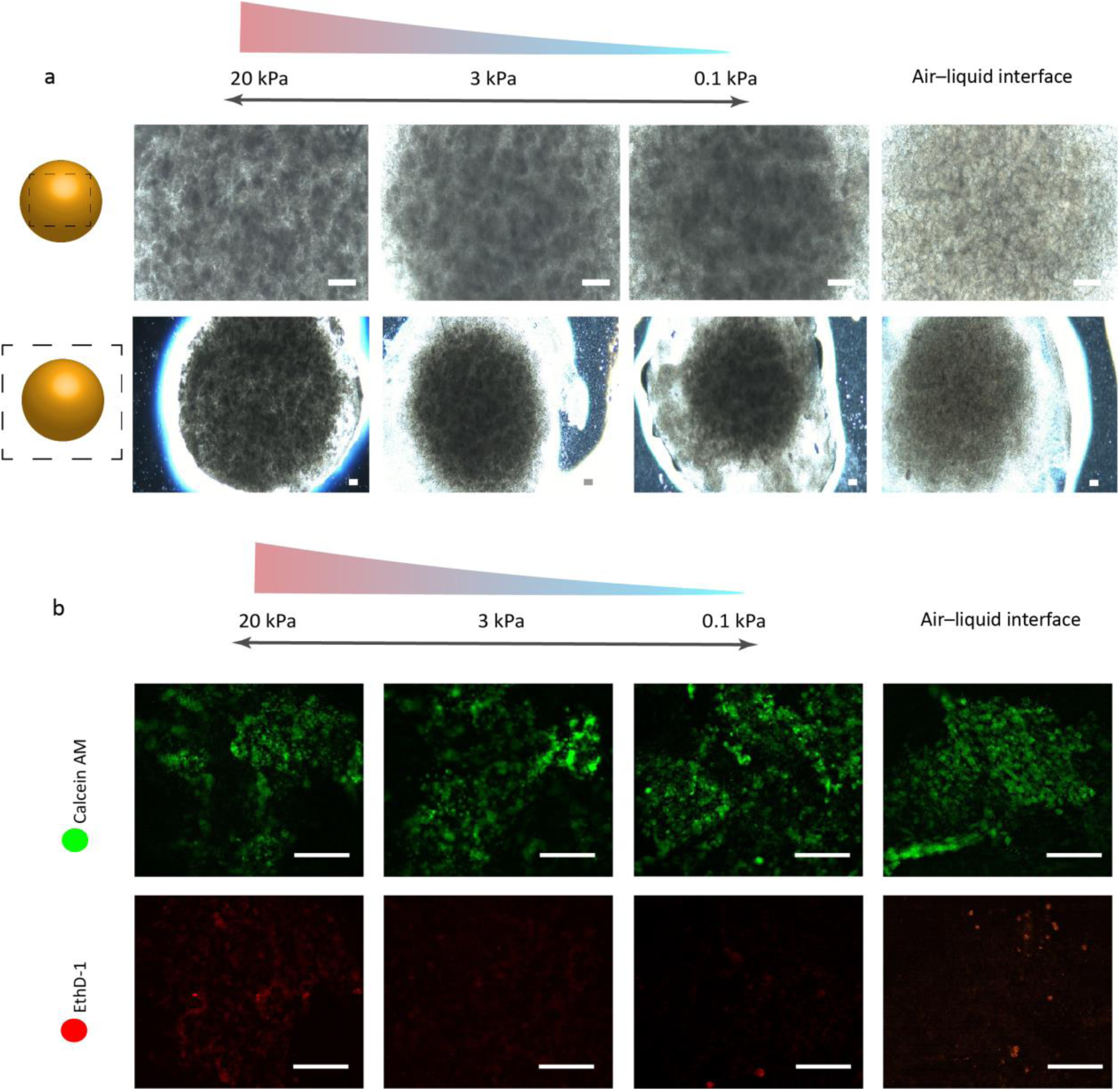
Organoid morphology and viability were maintained in the encapsulated hydrogels. a) Brightfield images of the organoids encapsulated in the different stiffness hydrogels or cultured at the air–liquid interface after 7+18 d. Schematic at the left indicates where images were taken. b) Live/dead assay with calcein AM (live) and EthD-1 (red) showed similar staining between the different culture environments. Scale bars: 100 µm. Representative images of N=3 organoid batches with n=3 organoids per batch.

**Figure S5.**
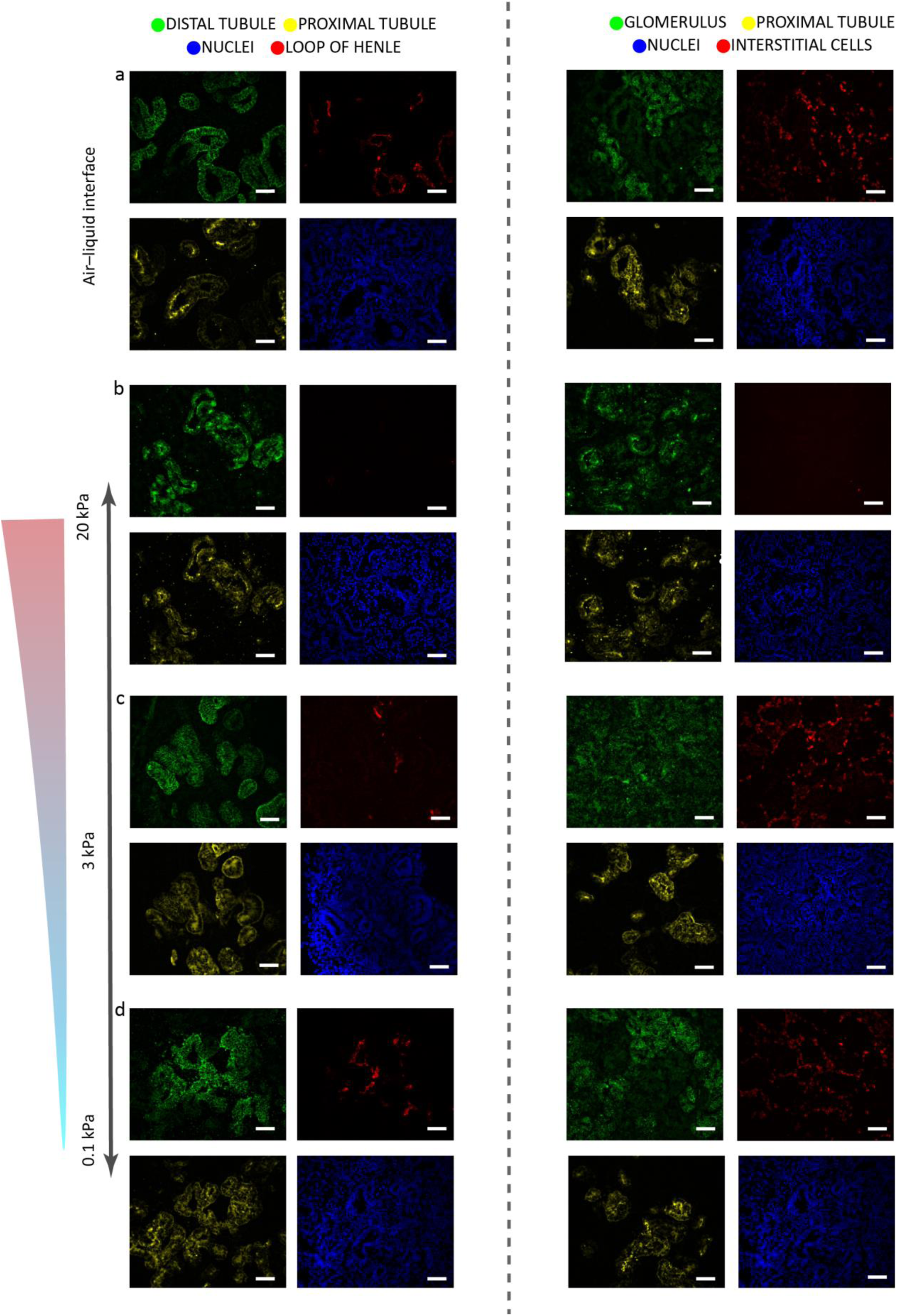
Single channels of immunohistochemistry in **Figure 2a-d** of glomeruli (nephrin: NPHS1, in green in third column from the left), proximal tubules (lotus tetragonolobus lectin: LTL, in yellow), loop of Henle (NKCC2 and SLC12A1, in red in second column from the left), distal tubules (E-cadherin: ECAD, in green in the far left column), and interstitial cells (homeobox protein Meis 1/2/3: MEIS1/2/3, in red in the far right column). DAPI staining (blue) for nuclei. Scale bars: 50 µm. Representative images of N=3 organoid batches with n=3 organoids per batch.

**Figure S6.**
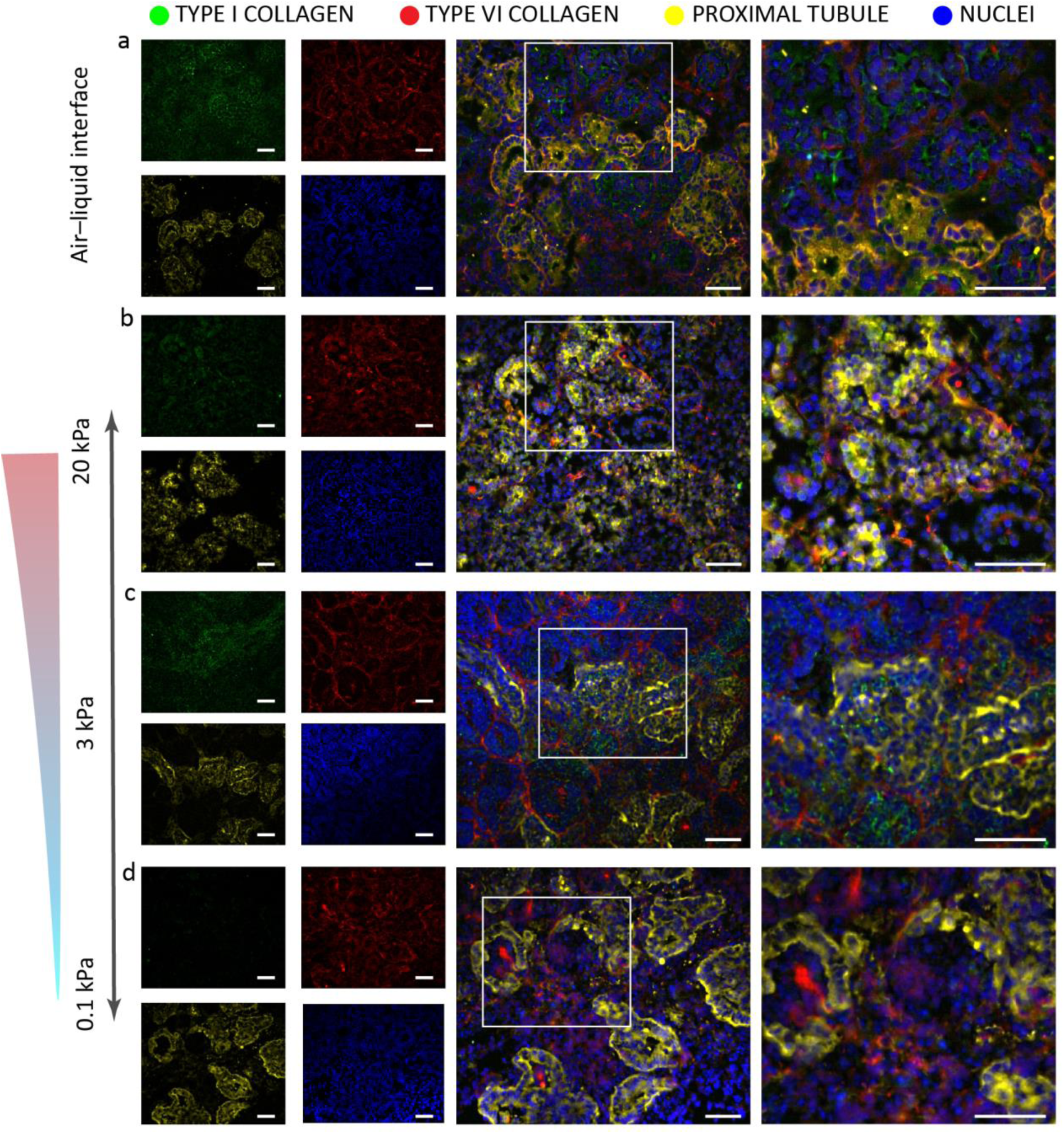
Reduced expression of collagen type 1a1 was observed in the encapsulated organoids. Immunohistochemistry for collagen type 1a1 (green), collagen type 6a1 (red), and LTL (yellow) on horizontally sectioned organoids encapsulated in the respective hydrogels (b–d) cultured on the air–liquid interface (a) at day 7+18. The reduced expression of type 1a1 collagen was observed in all encapsulated organoids (b–d) compared to organoids on the air– liquid interface (a). The white box denotes the area of interest enlarged in the respective right panel. Scale bars: 50 µm. Representative images of N=3 organoid batches with n=3 organoids per batch.

**Figure S7.**
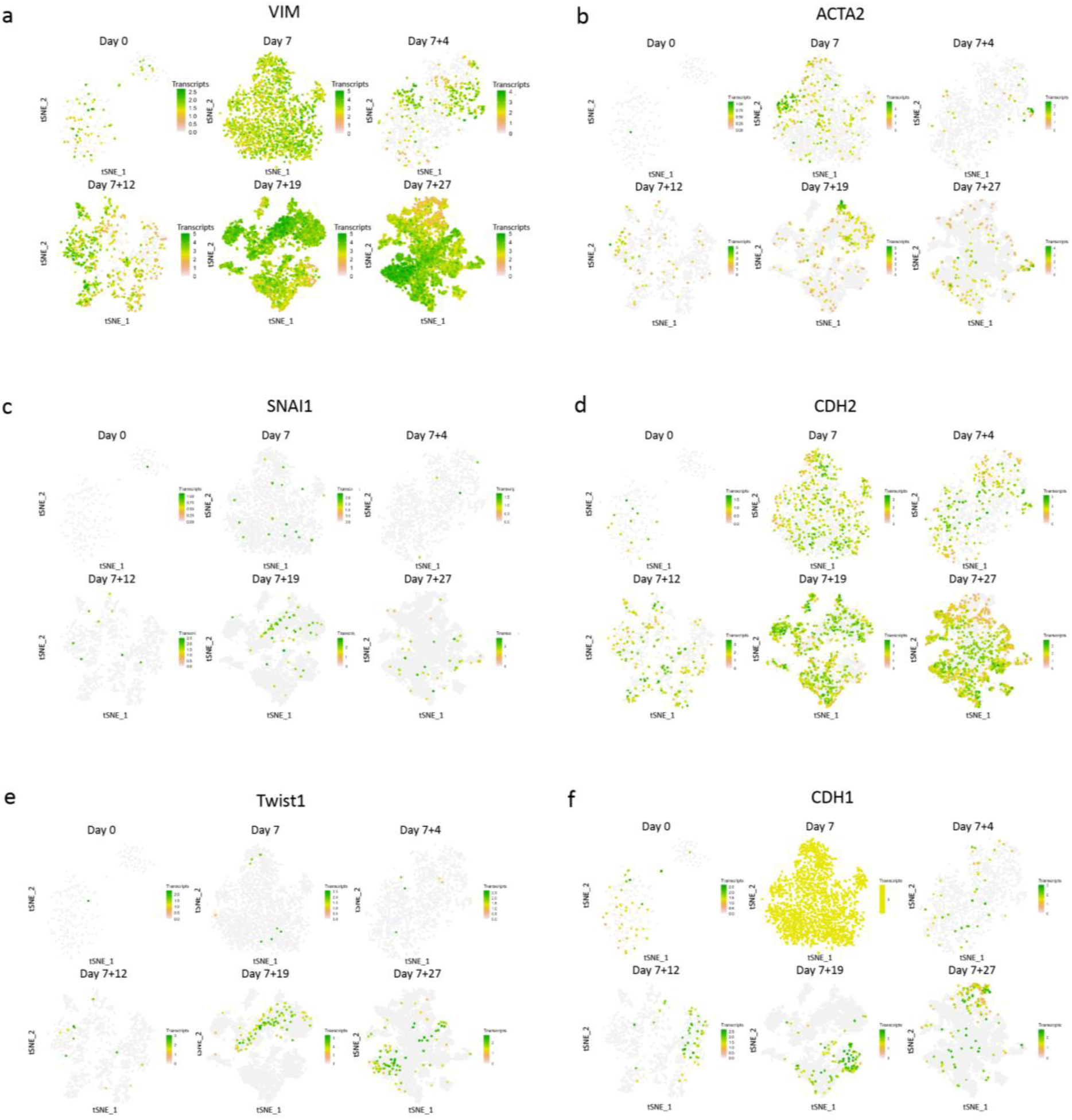
EMT-related gene expressions after prolonged air–liquid interface culture. Normalised gene expression of markers of interest shown in tSNE space of single-cell RNA (scRNA) sequencing data from the literature^25^. Cell populations expressing a) vimentin (VIM), b) actin alpha 2 smooth muscle (ACTA2), c) snail family transcriptional repressor 1 (SNAIL), d) N-cadherin (CDH2), and e) twist family bHLH transcription factor 1 (Twist1) increased in number when kidney organoids were cultured for 7+27 d. At the same time, cell populations expressing f) E-cadherin (CDH1) decreased during the culture.

**Figure S8.**
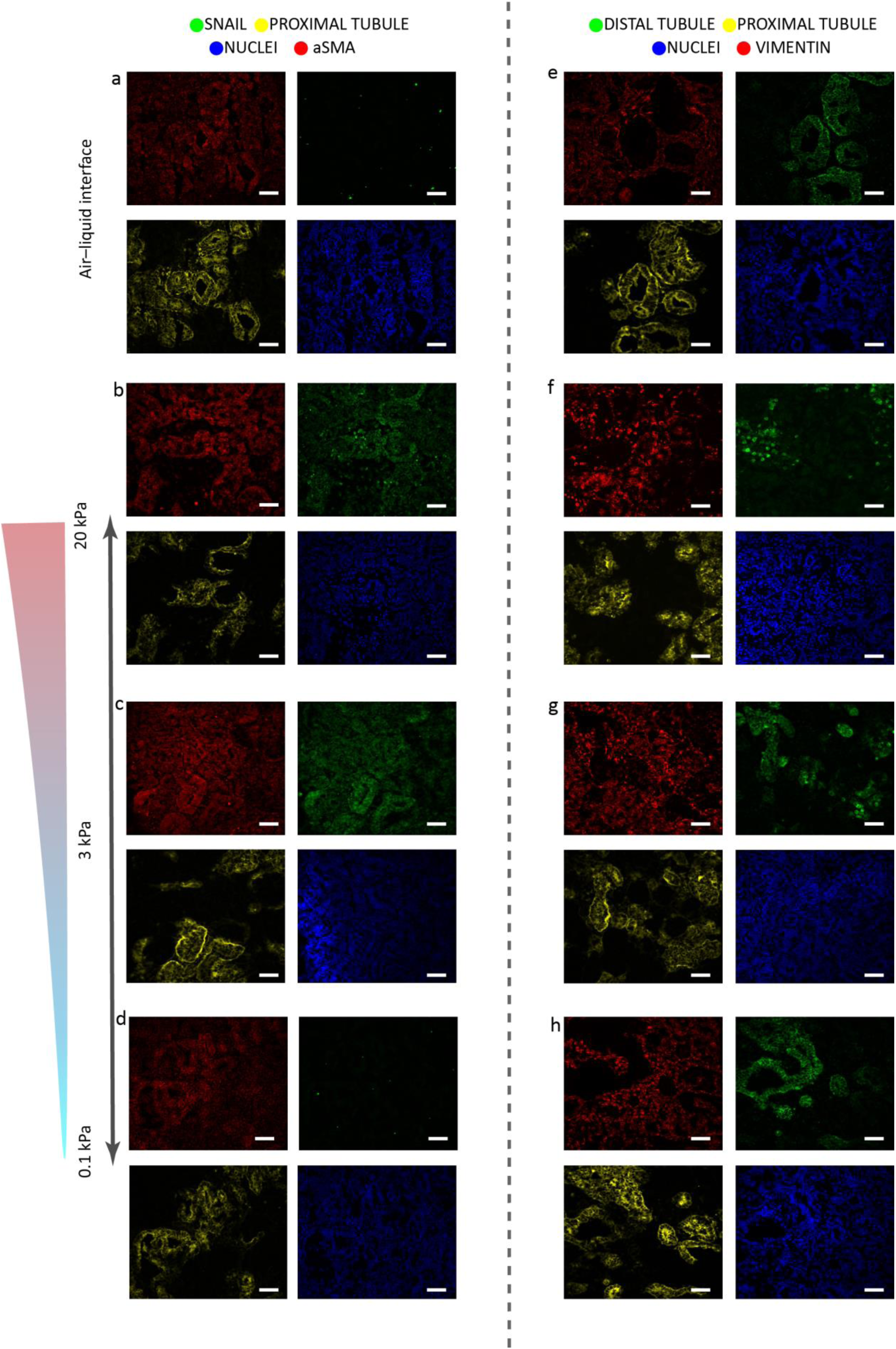
Single channels of immunohistochemistry in **Figure 3b-i** of proximal tubules (LTL, lotus tetragonolobus lectin, in yellow), distal tubules (E-cadherin: ECAD, in green in far right column), SNAIL (in green in second column from left), aSMA (in red, far left column), and vimentin (in red, third column from left). DAPI staining (blue) for nuclei. Scale bars: 50 µm. Representative images of N=3 organoid batches with n=3 organoids per batch.

**Figure S9.**
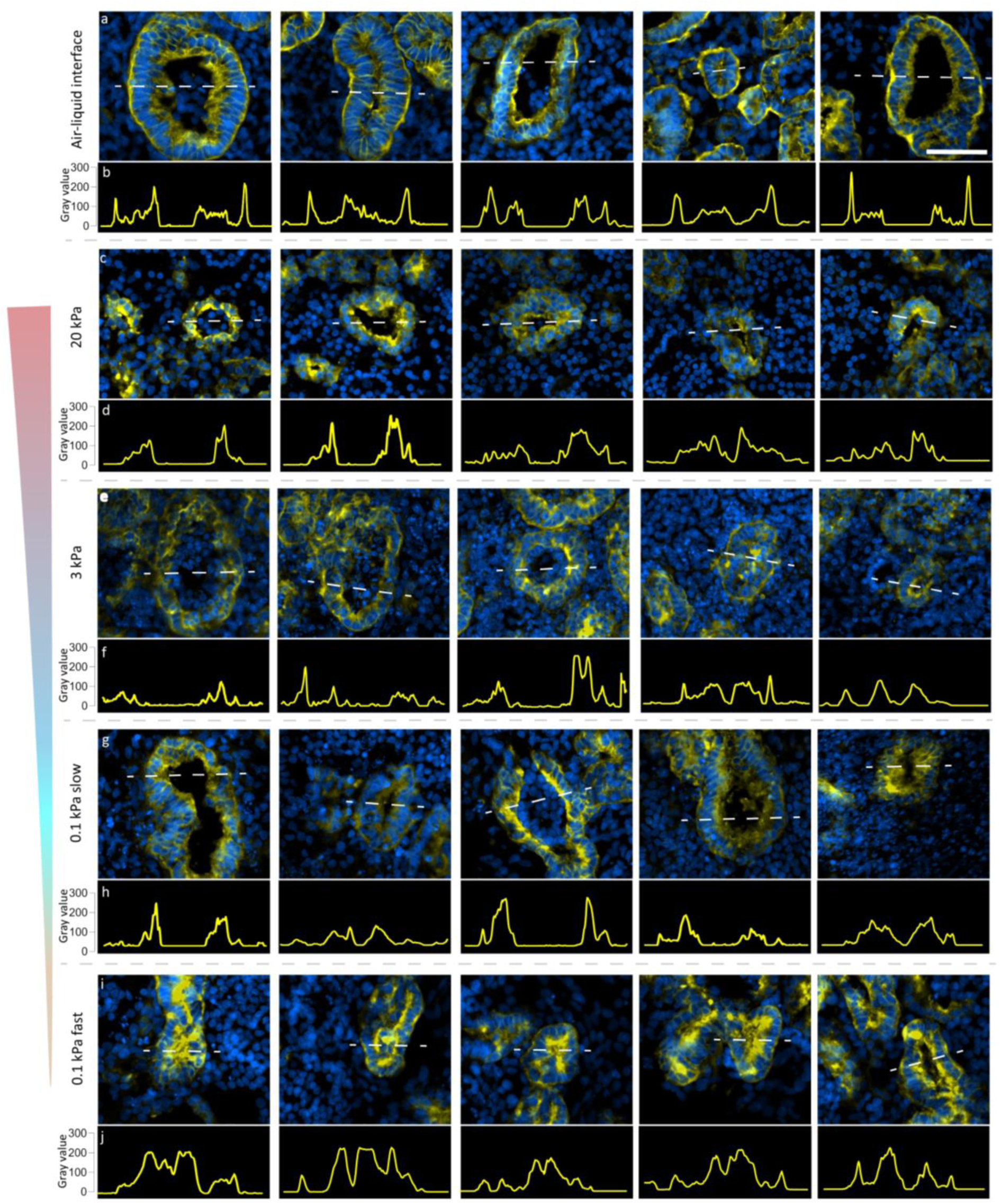
Lumen structures and apical enrichment. LTL staining was observed to be enriched in lumen structures of the organoids cultured in the 0.1 kPa hydrogels of fast (i-j) and slow (g-h) stress relaxing properties, compared to the air–liquid interface (a-b). While, smaller lumen structures and less LTL enrichment were observed for the 3 (e-f) and 20 (c-d) kPa hydrogels. Scale bar = 50 µm.

**Figure S10.**
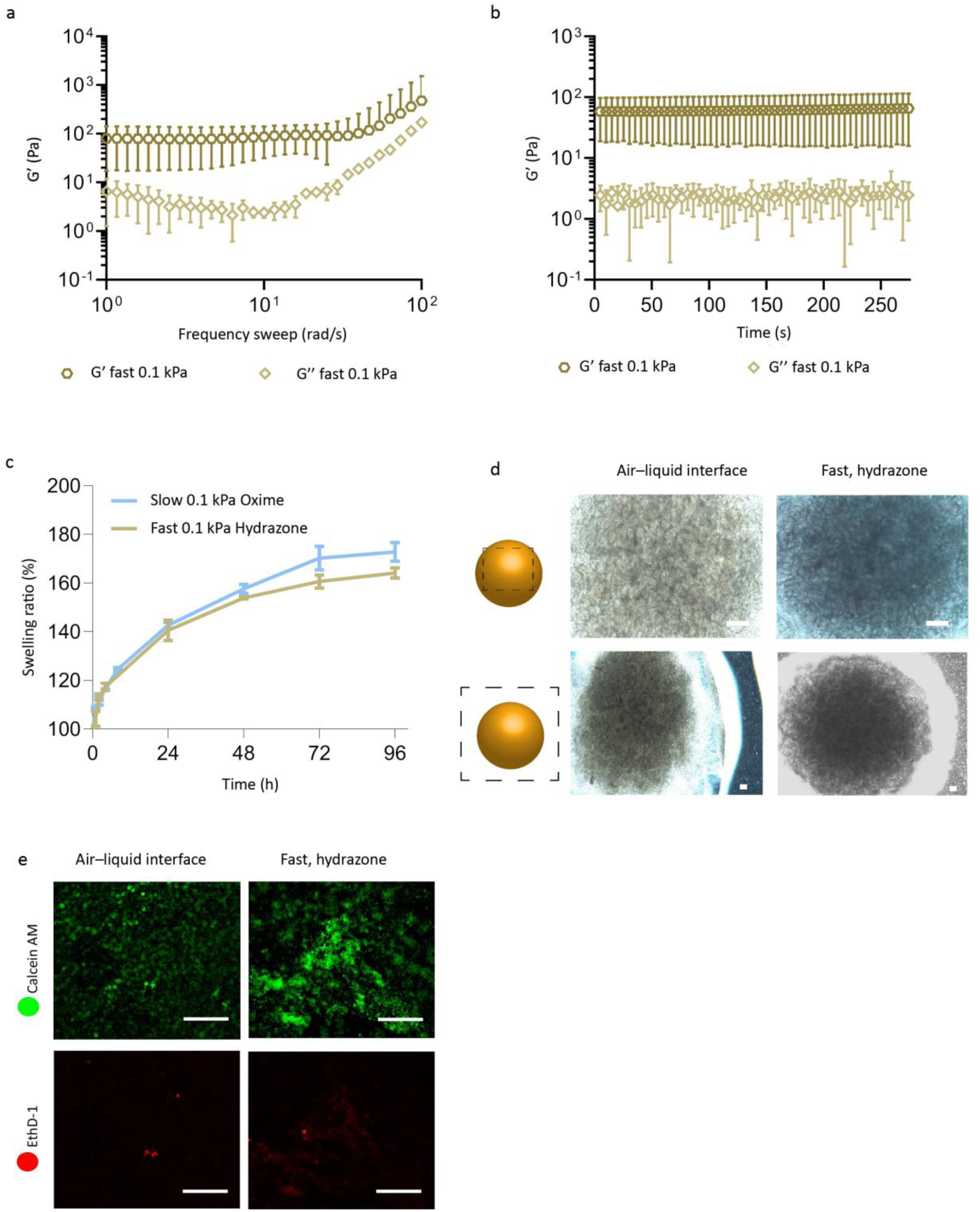
a) Frequency sweep data of the hydrogel (N=2) showed the hydrogel was not frequency dependent. b) Time sweep data of the hydrogel was used to determine the stiffness values of the hydrogel (N=2, Figure 1C). c) Swelling tests showed a swelling ratio of 165 % for the 0.1 kPa fast-relaxing hydrazone hydrogel after 96 h incubation, with no significant difference compared to the 172% swelling observed for the soft, slow-relaxing oxime cross-linked hydrogels (unpaired t-test, *p*=0.0662, N=3). d) Bright field images of organoids in the fast hydrazone hydrogel were unchanged compared to the air─liquid interface organoids. e) Live/dead assay with Calcein AM (live) and EthD-1 (red) showed no significant differences. Scale bars= 100 µm. Representative images of N=3 organoid batches with n=3 organoids per batch.

**Figure S11.**
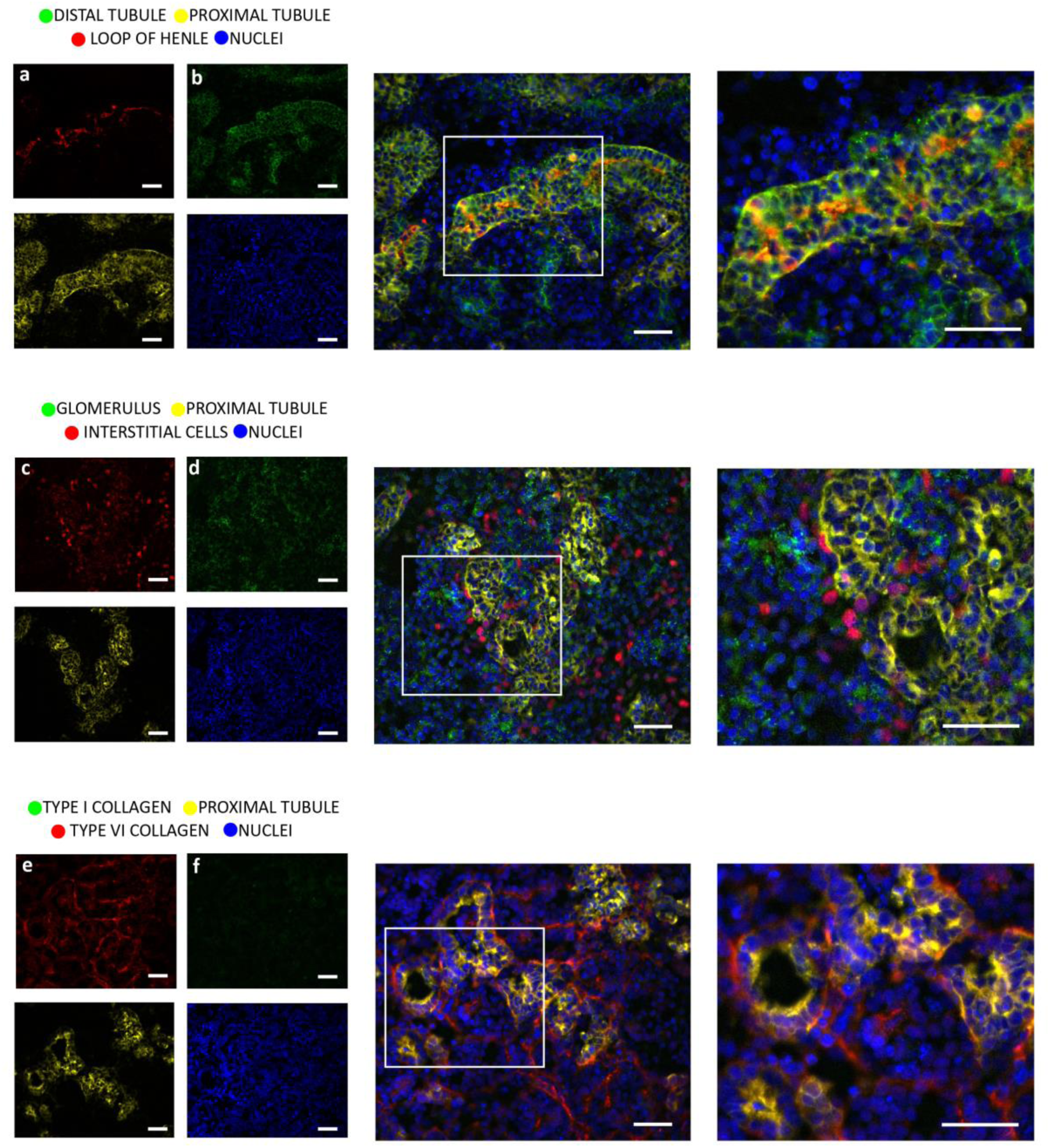
All renal cell types and reduced collagen type 1a1 were observed in the kidney organoids encapsulated in the 0.1 kPa, fast-relaxing hydrogel. Immunohistochemistry of kidney organoids encapsulated in the fast-relaxing hydrogel after 7+18 d. Staining was performed for the proximal tubules (LTL, lotus tetragonolobus lectin, in yellow), distal tubules (E-cadherin: ECAD, in green in a), loop of Henle (NKCC2: SLC12A1, in red in b), interstitial cells (homeobox protein Meis 1/2/3: MEIS1/2/3, in red in c), glomeruli (nephrin: NPHS1, in green in d), type 1a1 collagen (green in e), and type 6a1 collagen (red in f). DAPI staining (blue) for nuclei. Single channels are shown in the two left columns; merged images in the two right columns. The white box denotes the area of interest enlarged in the respective panel to the right. Scale bars: 50 µm. Representative images of N=3 organoid batches with n=3 organoids per batch.

**Figure S12.**
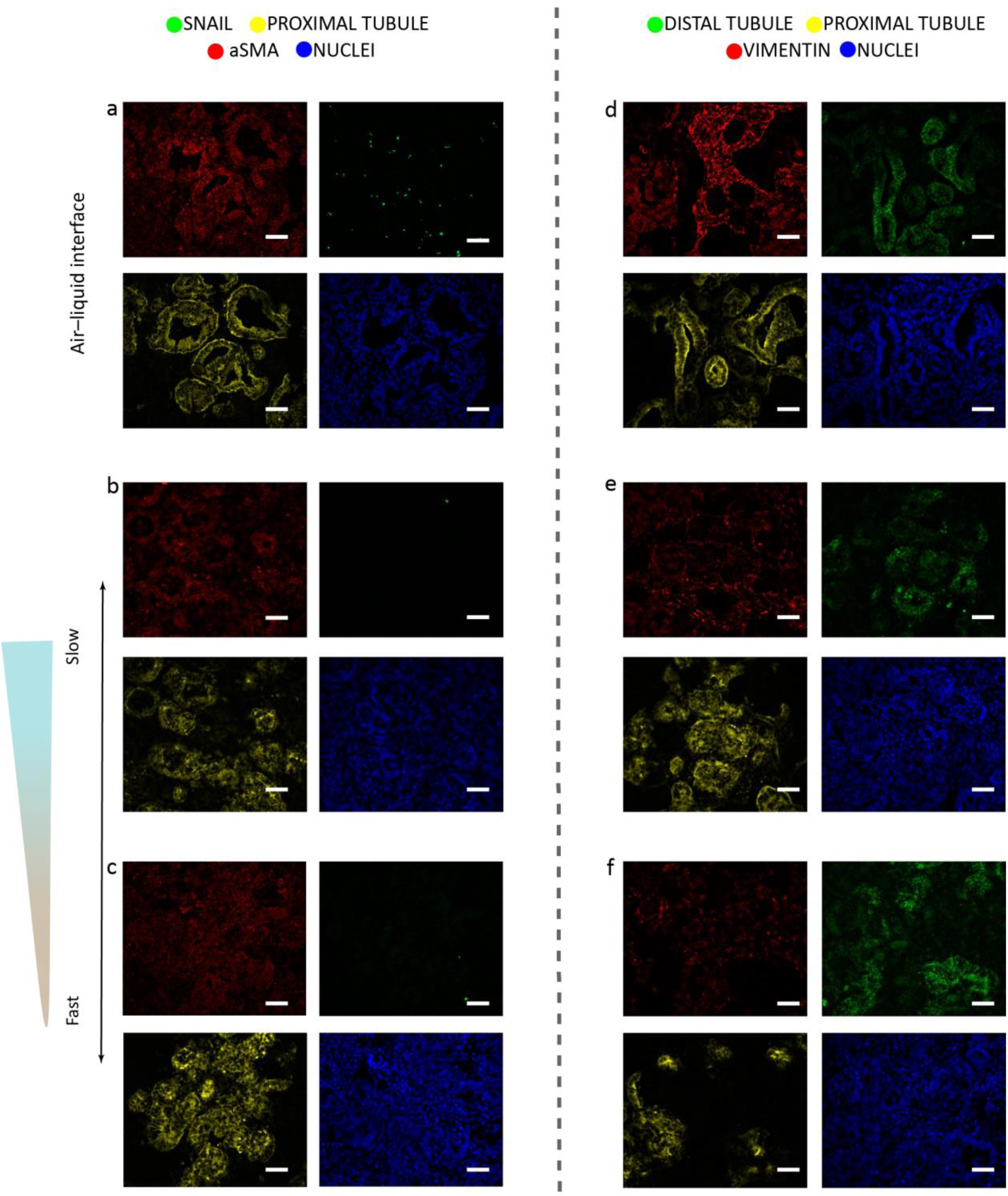
Single channels of immunohistochemistry in **Figure 4d-i** of proximal tubules (LTL, lotus tetragonolobus lectin, in yellow), distal tubules (E-cadherin: ECAD, in green in the far right column), SNAIL (green in the second column from the left), aSMA (red in the far left column), and vimentin (in red in the third column from left). DAPI staining (blue) for nuclei. Scale bars: 50 µm. Representative images of N=3 organoid batches with n=3 organoids per batch.

**Figure S13.**
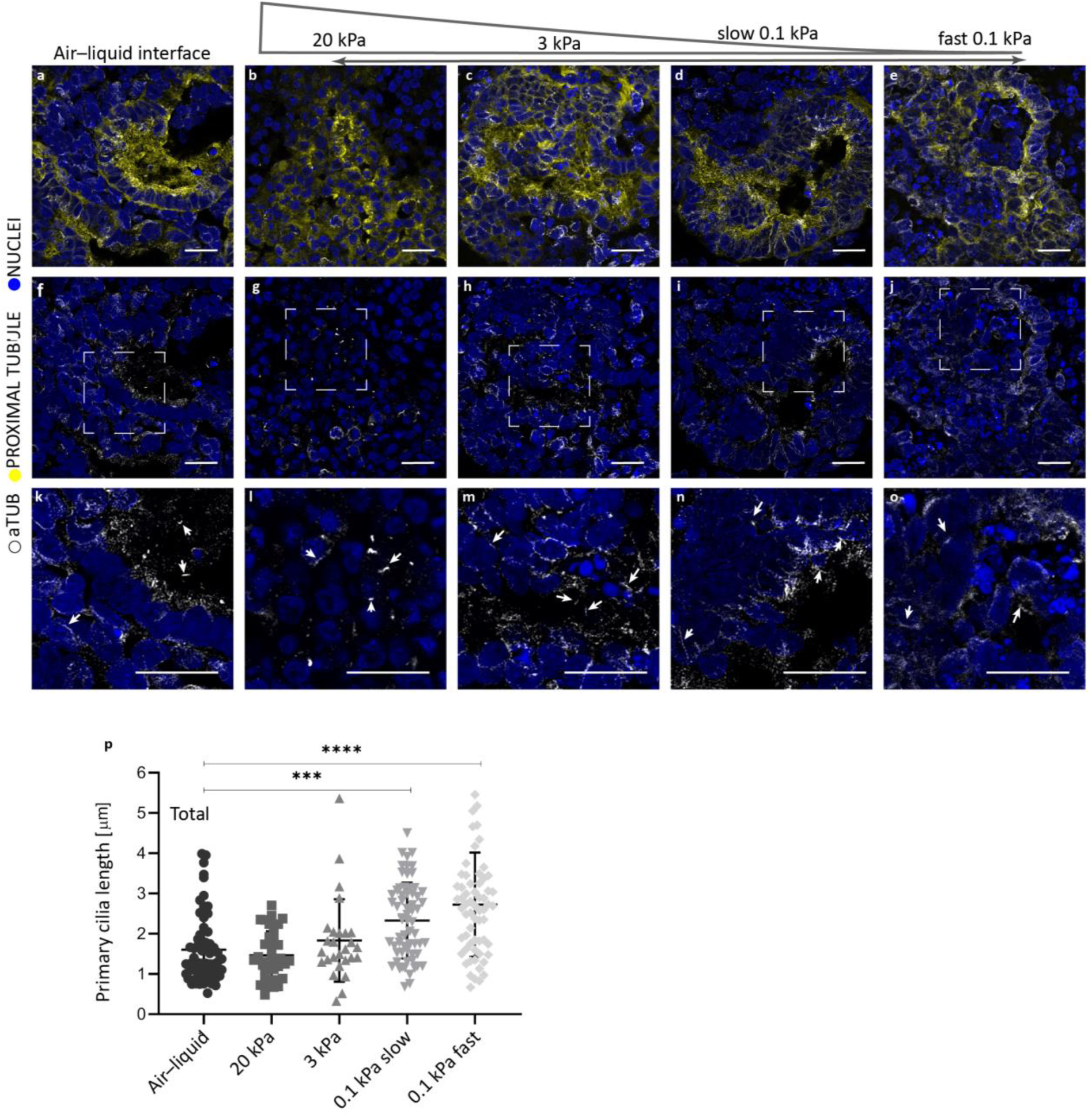
High resolution confocal images of primary cilia in a-f-k) air─liquid interface, b-g-l) 20 kPa, c-h-m) 3 kPa, d-i-n) 0.1 kPa slow-relaxation and e-j-o) 0.1 kPa fast relaxation. Scale bar= 20 µm. White boxes indicate area used for the zoomed in images k-o. Representative images of N=3 organoid batches with n=3 organoids per batch. p) Primary cilia length in the different hydrogels comparted to the air─liquid interface. Significantly, longer cilium were observed in the softer hydrogels. (***<0.005 and ***<0.0005, one-way ANOVA).

**Table S1.** Molecular weight averages by GPC. Number average (*M*_n_), weight average (*M*_w_), and dispersity (*M*_w_/*M*_n_) for pure alginate and two oxidised alginate (Oxi-Alg) batches used for the study.

| | $M_n$ [kDa] | $M_w$ [kDa] | $M_w/M_n$ [Đ] |
| --- | --- | --- | --- |
| **Pure Alg** | 130.7 | 316.2 | 2.46 |
| **Oxi-Alg1** | 28.3 | 69.6 | 2.46 |
| **Oxi-Alg2** | 83.5 | 185.3 | 2.21 |

**Table S2.** Volume, weights, and other properties of the prepared stock solutions and hydrogel solutions to form four different cross-linked alginate hydrogel systems used in these studies.

| Stock solutions |  |  |  |  |  |  |
| --- | --- | --- | --- | --- | --- | --- |
| Chemical | Weight [mg] | STEMdiff APEL2 or PBS [mL] |  | [mol/mL] | Wt% |  |
| Oxi-alg-1 or -2 | 165 | 2.75 |  | - | 6 |  |
| Oxime | 15 | 1.05 |  | 8.00×10<sup>-2</sup> | - |  |
| Hydrazone | 15 | 1.08 |  | 8.00×10<sup>-2</sup> | - |  |
| Hydrogel solutions and properties for rheometry |  |  |  |  |  |  |
| Hydrogel | Oxi-alg (-1 or -2) [μL] | Crosslinker [μL] | PBS [μL] | Crosslinker [μM] | Oxi-alg [% w] |  |
| 0.1 kPa | 51.7 (1) | Oxime, 3.9 | 99.4 | 2.02 | 2 |  |
| 3 kPa | 51.7 (1) | Oxime, 19.6 | 83.7 | 10.1 | 2 |  |
| 20 kPa | 103.3 (1) | Oxime, 39.2 | 12.5 | 20.2 | 4 |  |
| Fast 0.1 kPa | 51.7 (2) | Hydrazone, 19.6 | 83.7 | 10.1 | 2 |  |
| Hydrogel solutions and properties for organoid culture and swelling test |  |  |  |  |  |  |
| Hydrogel code | Oxi-alg (-1 or -2) [μL] | Crosslinker [μL] | STEMdiff APEL2 medium [μL] | Crosslinker [μM] | Oxi-alg [% w] | Stiffness [kPa] |
| 0.1 kPa | 166.7 (1) | Oxime, 12.6 | 320.7 | 2.02 | 2 | 0.08 |
| 3 kPa | 166.7 (1) | Oxime, 63.2 | 270.2 | 10.1 | 2 | 3 |
| 10 kPa | 333.3 (1) | Oxime, 126.3 | 40.4 | 20.2 | 4 | 20 |
| Fast 0.1 kPa | 166.7 (2) | Hydrazone, 63.2 | 320.7 | 10.2 | 2 | 0.1 |

